# FGF8-mediated gene regulation affects regional identity in human cerebral organoids

**DOI:** 10.1101/2023.12.22.572974

**Authors:** Michele Bertacchi, Gwendoline Maharaux, Agnès Loubat, Mathieu Jung, Michèle Studer

**Affiliations:** Univ. Côte d’Azur (UniCA), CNRS, Inserm, Institut de Biologie Valrose (iBV), Nice, France; GenomEast platform, Institut de Génétique et de Biologie Moléculaire et Cellulaire (IGBMC), Illkirch, France

**Keywords:** FGF8, multi-regional cerebral organoids, scRNAseq, dorsoventral and anteroposterior patterning, imbalance excitation-inhibition, spontaneous activity, neurodevelopmental disorders

## Abstract

The morphogen FGF8 establishes graded positional cues imparting regional cellular responses *via* modulation of early target genes. The roles of FGF signaling and its effector genes remain poorly characterized in human experimental models mimicking early fetal telencephalic development. We used hiPSC-derived cerebral organoids as an in vitro platform to challenge the effect of FGF8 signaling on neural identity and differentiation. We found that FGF8 treatment increases cellular heterogeneity leading to distinct telencephalic and mesencephalic-like domains that co-develop in multi-regional organoids. Within telencephalic domains, FGF8 affects the anteroposterior and dorsoventral identity of neural progenitors, the balance between GABAergic and glutamatergic neurons, thus impacting spontaneous neuronal network activity. Moreover, FGF8 efficiently modulates key regulators responsible for several human neurodevelopmental disorders. Overall, our results show that FGF8 signaling is directly involved in both regional patterning and cellular diversity in human cerebral organoids and in modulating genes associated with normal and pathological neural development.

## INTRODUCTION

A highly coordinated cascade of cellular and molecular events shapes the first phases of the embryonic development of the mammalian brain. This cascade involves distinct signaling pathways activated by gradients of diffusing morphogens, such as FGFs, BMPs, SHH and WNTs^1–5^. Brain morphogens instruct neural progenitors with spatial and temporal “coordinates” along the dorsoventral (D/V) and anteroposterior (A/P) brain axes and modulate early expression gradients of key developmental genes in a dose-dependent manner. These early stages become the basis for further key processes involved in brain development, such as progenitor proliferation, cell migration, neuronal differentiation, and circuit formation. Impairments of any of these mechanisms can result in multiple morphological abnormalities, ultimately leading to various neurodevelopmental disorders (NDDs).

Due to their pivotal roles in neural plate patterning^5^, FGF ligands and their receptors have been implicated in a plethora of brain malformations^6^ and disruption of the FGF pathways results in major cortical development defects^7–9^. Notably, despite a certain functional redundancy among FGF factors^6^, FGF8 is a major regulator of regional patterning and brain development. During early stages, FGF8 is crucial for the formation of the midbrain-hindbrain boundary^10,11^, a region called the isthmus and acting as a brain organizer, where FGF8 diffusing gradient controls the patterning of posterior brain regions^11–14^. During the formation of the telencephalic vesicles, FGF8 diffuses from the foremost anterior pole, a region called anterior neural ridge (ANR)^15,16^, forming a gradient that regulates proliferation, apoptosis, and identity acquisition along cortical axes^15,17–22^. Consistently, FGF8 diffusion is essential for the regulation of key telencephalic genes such as Foxg1^23^, a transcription factor required for regionalization and growth of the telencephalic vesicles. Moreover, FGF8 drives expansion of the cortex by activating the ERK signaling pathway, thus promoting self-renewal and pool increase of cortical radial glia cells^7^. FGFs function through binding to four highly conserved FGF receptors (FGFRs)^24^; among those, FGFR3 acts on brain size via regulation of progenitor proliferation and apoptosis^25^ and is specifically required for proper formation and regionalization of the occipitotemporal cortex^26,27^

Playing multiple functions during embryonic^5,28^ and adult age^6,24^, it is not surprising that FGF factors have been implicated in the onset of several NDDs^29^, and of adult psychiatric disorders, such as anxiety, depression, and schizophrenia^24,30^. FGF8 exerts its action *via* modulation of key developmental genes, which in turn control -directly or indirectly-important processes of neurogenesis, and whose genetic dysregulations can converge into pathological conditions. Nr2f1 is an example of developmental and pathogenic genes regulated by the FGF8 pathway. This nuclear receptor acts as a transcriptional regulator with multiple roles during mouse telencephalic development^31,32^. Mutations in NR2F1 lead to an emerging NDD, called Bosch-Boonstra-Schaaf optic atrophy syndrome (BBSOAS), in which patients have cognitive and visual impairments^33–36^.The anterior-most FGF8 telencephalic gradient negatively regulates mouse Nr2f1 during early development^37,38^ allowing its expression to follow an anterior-low to posterior-high gradient involved in the specification of areal identities in the dorsal telencephalon^39–44^. Other FGF8 downstream targets promote anterior identity by inhibiting Nr2f1^45,46^, building a cross-talk where FGFs and Nr2f1 operate opposite actions on the A/P cortical axis^31^. Interestingly, human NR2F1 is also expressed in a low anterior to high posterior expression gradient in the early telencephalon^47–49^, raising the possibility that FGFs could modulate similar downstream effector genes in the human brain. However, despite the great advances in the comprehension of FGF-related physiology and pathology, most studies use animal (often murine) models, in which depletion of FGF leads to severe phenotypes^50–53^. Hence, the role of FGFs and, in particular, FGF8 in human progenitor cells and neurons, remains poorly understood, as well as the contribution of FGF8 signaling and FGF8-target gene regulation in human diseases.

Self-organizing human brain organoids offer an unprecedented tool to model early steps of neural development *in vitro*^*54–62*^, bypassing obvious ethical and technical concerns when investigating early fetal brain development. Previous studies using 2D neural protocols^63–65^, followed by adaptations in 3D organoid protocols^66–68^, showed that pluripotent cells adopt a dorsal/anterior (*i*.*e*. telencephalic) neural fate when shielded from posteriorizing factors such as TGFβ, BMPs, retinoic acid, and WNTs. By building on previous methods^63,67,68^, we optimized a hybrid 2D/3D *in vitro* culture system for efficient formation of telencephalic FOXG1+ tissue and for assessment of FGF8-mediated effects on neural cell differentiation and identity acquisition. We found that FGF8-treated 3D organoids alter their regional identity when compared to non-treated ones, leading to the formation of multi-regional organoids with distinct co-developing brain domains. FGF8 can efficiently modulate NR2F1 and other brain-disease factors in human FOXG1+ telencephalic cells, pointing to a pivotal role of FGF8 signaling in the fine regulation of key neurodevelopmental and pathogenic pathways. Interestingly, FGF8 prolonged exposure affects cell identity along the D/V axis of the telencephalon together with an effect on A/P areal identity. As a result, FGF8-treated brain organoids have an imbalanced production of excitatory and inhibitory neurons and altered neural network formation. Our data redefine the role of FGF8 as a crucial morphogen for regional patterning and establishment of distinct D/V and A/P telencephalic identities in human cells, thus highlighting its importance in modulating the expression of key developmental and NDD-related genes during human brain organization.

### RESULTS

### An optimized 2D/3D organoid protocol promotes fast generation of telencephalic cells

Self-organizing human brain organoids allow to recapitulate and model early events of human fetal brain development *in vitro*^*54–56*^. Based on previous methods^63,67^, we optimized a human brain organoid protocol to study how FGF8 signaling impacts neuronal development and differentiation (**Figure 1A**). For fast and efficient induction of neural progenitors, we employed a dual inhibition of SMAD signaling paradigm^63^ by treating confluent 2D hiPSC cultures with TGFβ and BMP inhibitors (SB-431542 and LDN-193189, respectively). As endogenous WNT factors can inhibit the acquisition of an anterior fate^69,70^, a chemical WNT inhibitor (XAV-939) was added for optimal induction of telencephalic regional identity. Around day5-6, clusters of radially organized neural progenitors (*i*.*e*., neural rosettes) were visible in brightfield microscopy (Figure S1A). Consistently with the appearance of rosettes, analysis of key markers for pluripotency and neural differentiation by real-time RT-PCR showed efficient neural induction by day7 (**Figure 1B**), with down-regulation of stemness markers OCT4 and NANOG and upregulation of a molecular signature characteristic of antero-dorsal telencephalic neural progenitors (NPs) (SOX2+, PAX6+, SIX3+ and OTX2+). Notably, only NPs treated with XAV-939 (WNT inhibition; WNTi hereafter) efficiently upregulated the telencephalic marker FOXG1 at day10 compared to control samples (CTRL) (**Figure 1B**), as further confirmed by immunostaining (**Figure 1C,D**). To obtain 3D organoids, we dissociated day7 neural rosettes and re-aggregated early NPs as spherical aggregates (embryoid bodies, EBs; Figure S1A), which were included 24-hours later in Matrigel droplets. For optimal nutrient and oxygen distribution, EB-containing Matrigel droplets (named “cookies”) were cultured in miniaturized spinning bioreactors^67,68^. After few additional days of 3D culture, NPs spontaneously re-organized as multiple radially organized rosettes (**Figure 1E-F’**) and only WNTi organoids showed high FOXG1 expression compared to CTRL ones (**Figure 1E-I**). After 10-15 days (day35) of culture in neural differentiation medium (NDM), SOX2+ FOXG1+ NP rosettes started to be surrounded by differentiating neurons (**Figure 1J,J’**). From this stage onwards, the culture medium was supplemented with pro-survival and anti-apoptotic elements to supply optimal conditions for long-term cultures (long term survival medium, LTSM). At around day50, FOXG1+ NR2F1+ SOX2+ NP rosettes were surrounded by differentiated neurons (**Figure 1K-M’**), which were positive for neural-Tubulin (NTUB) staining (**Figure 1K,K’**) and expressed markers of cortical layers such as TBR1 and CTIP2 (**Figure 1M,M’**). In summary, we combined previous protocols for induction of human brain organoids, associating the high yield and rapidity of 2D neural induction with optimized growth of 3D neural structures in spinning bioreactors, in order to obtain anterior NPs in 7 days only and highly organized telencephalic organoids in 1 month-culture.

**Figure 1.**
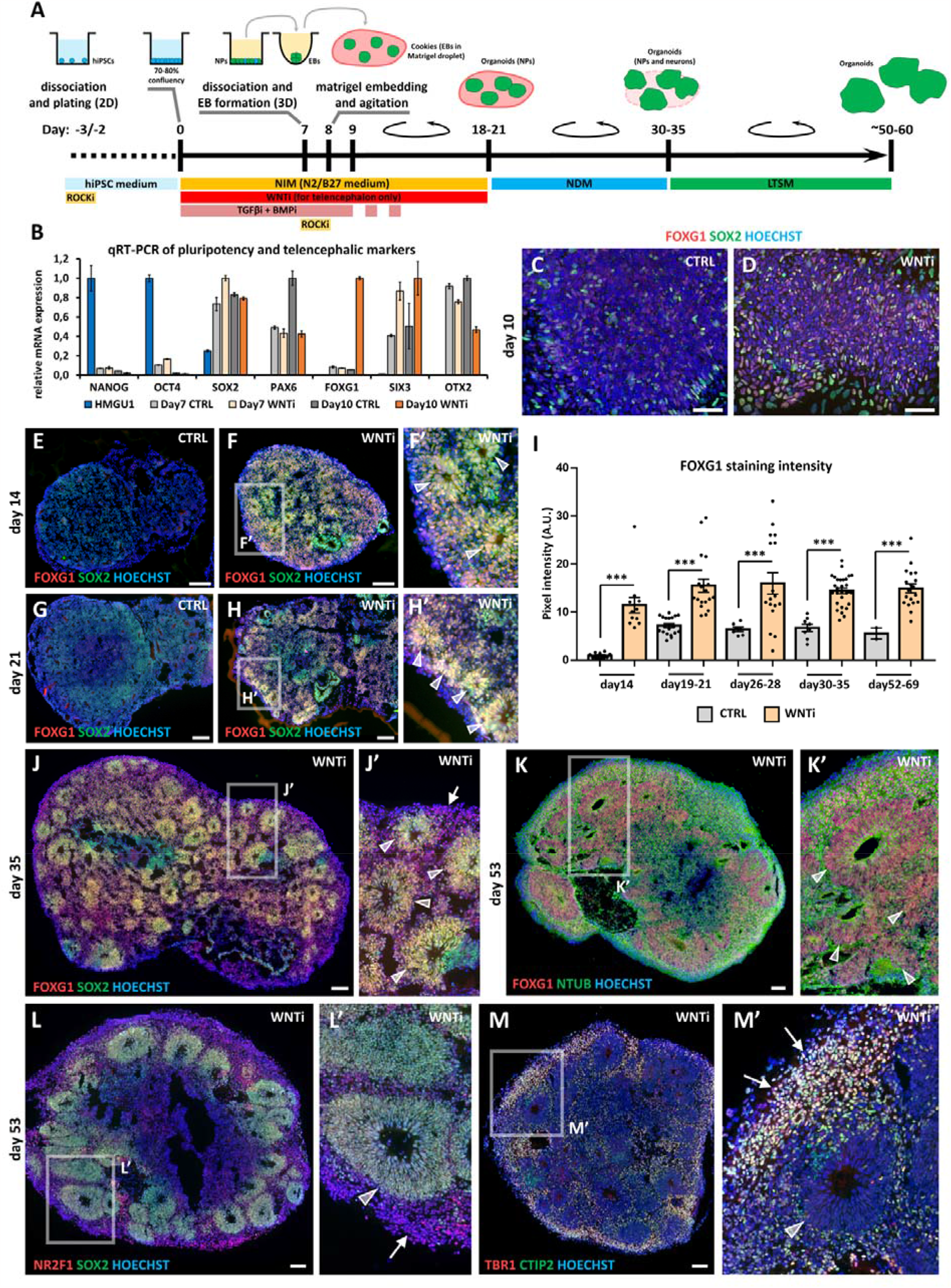
Hybrid 2D/3D protocol for fast formation of human cortical organoids. **A)** Schematic representation of the hybrid 2D/3D method for generating telencephalic/cortical human organoids in vitro by using triple TGFβ, BMP and WNT inhibition (SB-431542 5µM; LDN-193189 0.25µM; and XAV-939 2µM). At day7, cells were dissociated and re-aggregated in 96-well plates. (60.000 cells per aggregate). One day later, embryoid bodies (EBs) were included in Matrigel droplets (10 µl per droplet with 1-4 EBs). Cookies (matrigel droplets with EBs) were cultured in spinning bioreactors. **B)** Real time RT-PCR quantification of pluripotency (NANOG, OCT4) and telencephalic NP (SOX2, PAX6, FOXG1, SIX2 and OTX2) marker expression in undifferentiated HMGU1 hiPSCs and day7 or day10 control (CTRL) and WNT inhibited (WNTi) samples, as indicated. **C,D)** FOXG1 (red) and SOX2 (green) immunostaining of day10 2D neural cultures, in control condition (C) or after WNTi (D). **E-I)** FOXG1 (red) and SOX2 (green) immunostaining of day14 (E-F’) and day21 (G-H’) organoids, in CTRL or WNTi organoids as indicated. Graph in (I) shows pixel intensity quantification of FOXG1 immunostaining in CTRL and WNTi samples at different time points. White arrowheads in high magnification images point to NP neural rosettes. **J,J’)** FOXG1 (red) and SOX2 (green) immunostaining in day35 WNTi organoids. White arrowheads in high magnification images point to NP neural rosettes, while arrows highlight differentiating neurons surrounding the rosettes. **K-M’)** FOXG1 (red) and NTUB (green) (K,K’), NR2F1 (red) and SOX2 (green) (L,L’) and TBR1 (red) and CTIP2 (green) (M,M’) immunostainings in day53 WNTi organoids, highlighting FOXG1+ SOX2+ NR2F1+ NP rosettes/neuroepithelia (K-L’; indicated by white arrowheads in high magnification images) surrounded by TBR1+ CTIP2+ NR2F1+ differentiating cortical neurons (L’-M’; indicated by white arrows in high magnification images). Scale bars: 100 µm.

### FGF8 treatment modulates telencephalic target genes in brain organoids

In the developing mouse brain, distinct sources of diffusing FGF8 fine-tune the expression of several genes. In WNTi organoids, we found that early FGF8 treatment (starting at day5; Figure S1B) reduced FOXG1 expression, in accordance with early FGF8 inducing posterior and not anterior identity^30^. The addition of FGF8 to the culture medium starting at day10-11 (hereafter, WNTi+FGF8 condition) (**Figure 2A**) did not affect FOXG1 expression (**Figure S1B,C**), hence we chose day10-11 as a starting time point for FGF8 treatment. Real-time RT-PCR analysis of known FGF8 target genes (SPRY4, DUSP6, ETV4, ETV1 and ETV5) confirmed that our set-up efficiently activated FGF signaling in day20 and day30 organoids (**Figure 2B**). As a first read-out of FGF8 treatment on the expression of target telencephalic genes, we stained control (WNTi) and treated (WNTi+FGF8) organoids for NR2F1 and FOXG1 at different time points. NR2F1 expression, still low at day19 but higher at day26, was clearly modulated by FGF8 (**Figure S2A-H’**). Real-time RT-PCR confirmed FGF8-mediated inhibition of NR2F1 at day20 and day30 (**Figure 2C**), while FOXG1 levels were partially affected upon FGF8 treatment at day30 (**Figure 2C**). Despite this, WNTi+FGF8 organoids still largely expressed FOXG1 at later stages (day35 and day53) (**Figures 2D-F** and **S2I-J’**) together with efficient modulation of NR2F1 levels (**Figures 2G-I** and **S2K-L’**). FGF8 was maintained in culture medium until day ∼60 to prevent an early increase of NR2F1 expression back to normal levels (**Figure S2M-P’**). Thus, FGF8 was efficient in modulating its target gene NR2F1 in telencephalic organoids starting at day19-21 and until day69-74 while maintaining FOXG1 expression at high levels (**Figure S2Q,R**), demonstrating the efficacy of FGF8 treatment *in vitro* and the evolutionary conservation of the NR2F1-FGF8 regulatory molecular axis from mice to humans.

**Figure 2.**
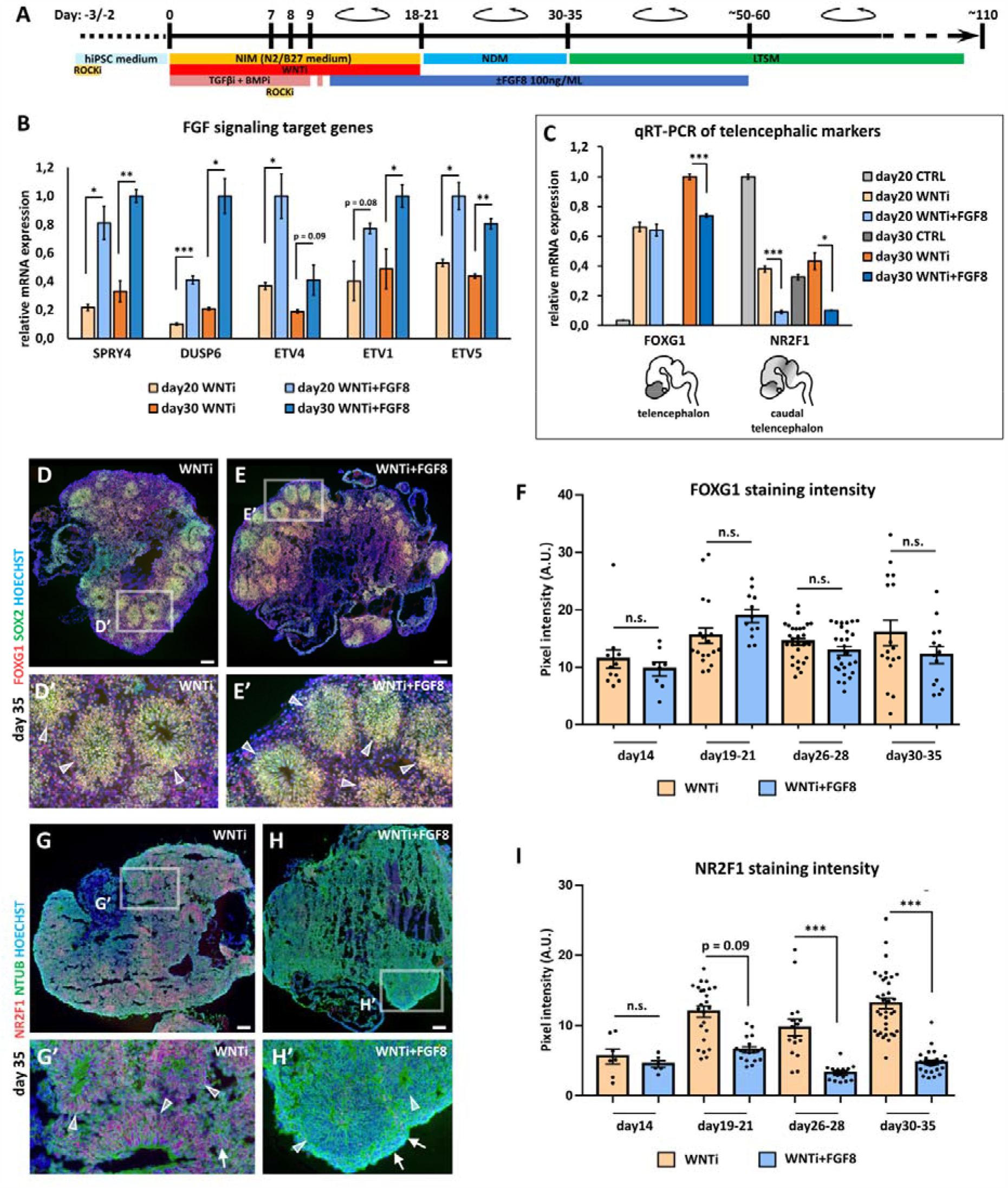
FGF8-mediated control of target gene expression in FOXG1+ telencephalic organoids. **A)** Schematic representation of the hybrid 2D/3D method for performing FGF8 treatment on telencephalic/cortical human organoids in vitro. FGF8 (100ng/ml) was added to neural progenitor patterning medium (NPPM) starting at day10-11 (blue bar) and kept for following culture steps until day ~50-60. **B)** Real time RT-PCR of the expression levels of FGF8-target genes (SPRY4, DUSP6, ETV4, ETV1 and ETV5) in day20 or day30 WNT inhibited (WNTi) or FGF8-treated (WNTi+FGF8) organoids, as indicated. **C)** Real time RT-PCR quantification of the expression of FOXG1 (telencephalic marker) and of NR2F1 (FGF8 target gene and caudal telencephalic marker) in day20 or day30 control (CTRL), WNT inhibited (WNTi) and FGF8 treated (WNTi+FGF8) samples, as indicated. FGF8 treatment efficiently down-regulated NR2F1 levels in WNTi+FGF8 organoids, compared with WNTi ones. **D-F)** FOXG1 and SOX2 (red and green, respectively, in D-E’) immunostainings on day35 WNTi and WNTi+FGF8 organoids, as indicated. FGF8 treatment did not significantly interfere with FOXG1 expression. White arrowheads in high magnification images point to double SOX2 NR2F1 NPs in neural rosettes. Graph in F shows pixel intensity quantification of FOXG1 immunostaining in WNTi and WNTi+FGF8 organoids at different time points, as indicated. G-I) NR2F1 and NTUB (red and green, respectively, in G-H’) immunostainings on day35 WNTi and WNTi+FGF8 organoids, as indicated. FGF8 treatment efficiently modulated NR2F1 expression (compare G with H). High magnifications in G’ and H’ show neural rosettes (NTUB^low^; white arrowheads) and differentiating neurons (NTUB^high^; white arrows), both expressing NR2F1 (red) in WNTi organoids but lacking NR2F1 in WNTi+FGF8 ones. Graph in I shows pixel intensity quantification of NR2F1 immunostaining in WNTi and WNTi+FGF8 organoids at different time points, as indicated. Scale bars: 100 µm.

### Single-cell RNA sequencing (scRNAseq) unveils multiple progenitor and neuronal classes in human organoids

To investigate the transcriptomic signature of cerebral organoids upon FGF8 treatment, we employed a scRNAseq approach to compare control (WNTi) with FGF8-treated telencephalic organoids (WNTi+FGF8). Two distinct batches of both WNTi and WNTi+FGF8 organoids showed high reproducibility in terms of cell cluster compositions (**Figure 3A**) allowing us to pool together individual batches to reach a higher number of cells per cluster (n° of cells per cluster in **Figure S3A**). Bioinformatic analysis of the whole cell population (WNTi and WNTi+FGF8 cells together) identified 15 distinct cellular clusters (shown as UMAP projections; **Figure 3B**), whose composition was recognized *via* the expression of well-known reference markers as a read-out of cell identity (**Figures 3C** and **S3B**). Cell clusters comprised NPs (NESTIN+, SOX2+ and HES5+ cells in clusters 2/5/8/9/12/13/14/15), some of which were actively proliferating (KI67+ and TOP2A+), and neurons (MAPT+ cells in clusters 1/3/4/6/7/10/11). Expression of neurogenic (NEUROD4 and NEUROG1) and post-mitotic neuronal markers (DCX, NEUN and MAPT) highlighted the existence of both differentiating and differentiated neurons (**Figures 3C** and **S3B**). EOMES (also known as TBR2) expression identified intermediate progenitors, while expression of HOPX, TNC and FAM107A pointed to both apical and basal radial glia^71^. Interestingly, a *bona fide* marker for late truncated radial glia (CRYAB)^72^, which is normally expressed in the neocortex *in vivo* starting at GW16.5, was specifically expressed in NP cluster 13 (**Figure 3C**). We reasoned that multiple NP and neuronal types co-existed in telencephalic organoids, and trajectory analysis confirmed that NP clusters (clusters 2/5/8/9/12/15) gradually converted into post-mitotic neurons (clusters 1/3/4/6/7) (**Figure 3D**), suggesting that key steps of human brain development were recapitulated *in vitro*.

**Figure 3.**
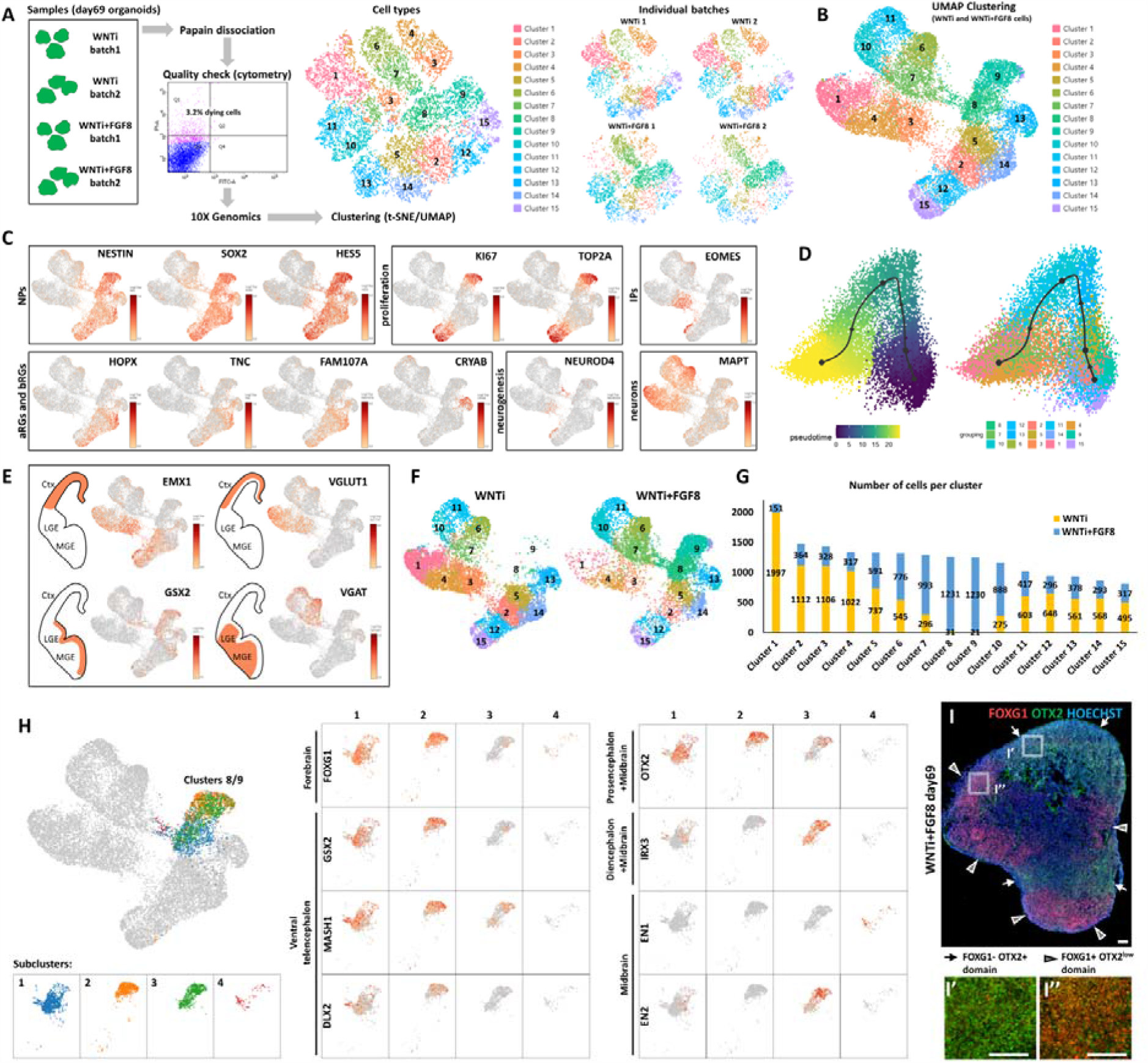
Single-cell RNA sequencing (scRNAseq) analysis highlighting FGF8-induced cellular and molecular changes in human organoids. **A)** Schematic representation of the experimental set-up for scRNAseq of control (WNTi) and FGF8-treated (WNTi+FGF8) telencephalic organoids at day69. Two independent batches of WNTi and of WNTi+FGF8 organoids (each containing 2 to 3 organoids) were dissociated into single cells and processed with Chromium (10X Genomics technology). Identified cells were clustered and represented in a 2D-space using t-SNE and UMAP algorithms. **B)** UMAP clustering of WNTi and WNTi+FGF8 cells, showing 15 distinct clusters. **C)** Expression level of known markers identifying neural progenitor cells (NPs; NESTIN, SOX2 and HES5), proliferating progenitors (KI67 and TOP2A), intermediate progenitors (IPs; EOMES), apical and basal radial glia cells (aRGs and bRGs; HOPX, TNC, FAM107A and CRYAB) and differentiating or differentiated neurons (NEUROD4 and MAPT, respectively). **D)** Trajectory analysis evaluating the most probable developmental trajectory linking clusters of all cells, from NP clusters (2/5/8/9/12/15) to post-mitotic cell types (notably clusters 1/3/4/6/7). **E)** Expression level and cluster distribution of the glutamatergic dorsal markers EMX1 and VGLUT1, and of the GABAergic ventral markers GSX2 and VGAT, suggesting coexistence of both glutamatergic and GABAergic NPs and neurons in FOXG1+ telencephalic organoids. **F,G)** UMAP clustering of WNTi and WNTi+FGF8 cells separately, showing 15 distinct clusters and the abundance of each cluster in the two experimental conditions. For each cluster, the number of cells originated from control WNTi organoids (yellow) or from treated WNTi+FGF8 organoids (blue) is indicated in G. **H)** UMAP projection of day69 organoid scRNAseq data showing 4 cellular groups (left), as identified by sub-clustering analysis on WNTi+FGF8 clusters 8 and 9. Panels (centre and right) show the expression levels of forebrain (FOXG1), ventral telencephalon (GSX2, MASH1, DLX2), forebrain/midbrain (OTX2), diencephalon/mesencephalon (IRX3) and mesencephalon markers (EN1, EN2) in the 4 sub-clusters. **I-I’’)** FOXG1 (red) and OTX2 (green) immunostaining in day69 treated (WNTi+FGF8) organoids, showing the presence of segregated FOXG1+ and FOXG1-domains. White arrows point to FOXG1-OTX2+ non-telencephalic regions (high magnification in I’), while arrowheads highlight FOXG1+ OTX2^low^ telencephalic region (high magnification in I’’). Abbreviations: Ctx, cortex; MGE, medial ganglionic eminence; LGE, lateral ganglionic eminence.

To assign the identity of distinct cell clusters in a more unbiased way, we used the VoxHunt spatial brain mapping and heatmap of similarity score tools^73^ (**Figure S3C-E**), which evaluate the similarity between the expression profile of each cluster and that of spatial and single-cell transcriptome reference datasets. We established high resemblance of most clusters to the dorsal telencephalon (pallium), with clusters 2/5/12/14/15 showing a high similarity score with progenitors, and clusters 1/3/4 including cortical neurons (**Figure S3C-E**). However, some ventral telencephalic (subpallium) clusters were also present (clusters 6 and 7), in line with previous reports showing that ventral GABAergic identity can spontaneously arise in dorsally patterned organoids^74^. Notably, clusters 8, 9, 10 and 11 showed a mixed identity, as they scored high with the dorsal but also with the ventral telencephalon and/or with more posterior mesencephalic regions (**Figure S3E**). Analysis of known glutamatergic and GABAergic markers substantiated the D/V identity of organoid NPs and neurons, that could be subdivided into two pools of glutamatergic (EMX1+ and VGLUT1+) and GABAergic (GSX2+ and VGAT+) cells (**Figure 3E**). Additionally, trajectory analysis of different clusters highlighted a very dynamic state of developmental trajectories (**Figure S3F**). Together, transcriptomics data point to a mixed dorsal (glutamatergic) and ventral (GABAergic) identity in our organoid model, which after 2 months of culture seems to display the cellular and molecular signature of GW16.5 human neocortex.

### FGF8 treatment increases brain regional heterogeneity

While widespread FOXG1 expression supported the general telencephalic identity of both WNTi and WNTi+FGF8 treated organoids (**Figures S4A,B**), qRT-PCR showed a partial reduction of the anterior marker SIX3, induction of OTX2 (a telencephalic marker at early stages^75^ before becoming restricted to more posterior regions at later stages^76^), and induction of the mesencephalic marker EN2 in WNTi-FGF8 organoids (**Figure S4C**), suggesting that long-term FGF8 treatment might induce other regions than the forebrain. Hence, we explored the expression of key markers of A/P regional identity by focusing on clusters 8 and 9, as these cellular populations were almost absent in control (WNTi) organoids and only appeared in treated (WNTi+FGF8) samples (**Figures 3F,G** and **S4**). Although cluster 8/9 cells were largely positive for telencephalic markers such as FOXG1 (50% positive cells) and SFRP1 (70% positive cells) (**Figure S4A,B,D,E**), they also displayed expression of the diencephalic gene SIX3 (>40% positive cells) and mesencephalic markers OTX2 (>30% positive cells), IRX3 (20% positive cells) and EN2 (10% positive cells) upon FGF8-treatment (**Figure S4D,E**). These data indicate that clusters 8/9, which are only present in FGF8-treated organoids, are mainly composed of telencephalic progenitors (FOXG1+, SFRP1+, GAS1+ and FZD8+) but also contain some diencephalic (SIX3+) and mesencephalic (IRX3+, OTX2+, EN2+) cells (**Figure S4E**), suggesting concomitant FGF8-driven induction of non-telencephalic regional identities.

To further distinguish cell types in scRNAseq clusters 8 and 9, we performed a sub-clustering analysis, which detected four main cellular groups (**Figure 3H**). While two sub-clusters expressed FOXG1 together with ventral telencephalic markers (GSX2, DLX2 and MASH1), the remaining two were negative for FOXG1 but positive for OTX2, IRX3, EN1 and/or EN2 (**Figure 3H**). To directly visualize the co-existence of distinct organoid domains, we performed a double staining for FOXG1 (telencephalon) and OTX2 (diencephalon/mesencephalon) on treated (WNTi+FGF8) organoids (**Figure 3I-I’’**) and confirmed that FOXG1-negative domains were indeed positive for OTX2, most probably corresponding to the diencephalic/mesencephalic sub-clusters identified in scRNAseq data. Collectively, we show that while WNTi organoids developed into uniform FOXG1+ telencephalic organoids mostly expressing cortical markers, FGF8-treated organoids showed co-existing and segregated regional domains. Thus, high levels of FGF8 can increase the complexity of human cultured organoid *in vitro*, by inducing the formation of multi-regional organoids in which different brain regions co-exist and co-develop in the same brain aggregate.

### FGF8 alters dorso/ventral cell specification and network neuronal activity of telencephalic domains

While the expression of diencephalic/mesencephalic markers was limited to clusters 8/9 only (**Figure S4D,E**), most of the remaining clusters expressed high levels of FOXG1 (**Figure S4A,B**). Hence, we focused on the effect of FGF8 treatment on these more abundant FOXG1+ telencephalic populations. By comparing control (WNTi) *versus* treated (WNTi+FGF8) samples in terms of cell abundance per cluster, we noticed an increase in cell number in clusters 6/7 -corresponding to ventral cells-upon FGF8 treatment (**Figures 3F,G**). On the contrary, clusters corresponding to glutamatergic NPs (clusters 2/5/12/14/15) and neurons (clusters 1/3/4) were abundantly populated in control organoids but contained fewer cells in FGF8-treated ones (**Figure 3F,G**). As the different abundance of cell clusters in control or treated organoids implied a different cellular composition upon FGF8 treatment, we explored the expression of key dorsal glutamatergic or ventral GABAergic markers on WNTi and WNTi+FGF8 UMAP projections (**Figure 4A,B**). Glutamatergic NP and neuronal markers such as EMX1, NEUROD6, NEUROD2, TBR1, SOX5, CTIP2, LHX2, NEUROG2, NF1A and VGLUT1 were highly reduced in FGF8-treated organoids (**Figure 4A** and S5A), consistently with a lower cell number in clusters 1/3/4. Notably, the upper cortical layer marker SATB2 was completely absent in WNTi+FGF8 cluster 1 (**Figure S5A**). On the contrary, ventral GABAergic or striatal markers such as MASH1, DLX1, DLX2, PBX3, GAD1 and GAD2 were increased (**Figure 4B** and S5B) in clusters 6/7/8/9, supporting a ventralization of FGF8-treated samples compared to non-treated ones. An unbiased VoxHunt analysis of the correlation score with brain areas confirmed that FGF8 treatment decreased dorsal (pallial), while increasing ventral (subpallial) telencephalic properties (**Figure S5C**). Additionally, FGF8 slightly increased the similarity score with more posterior areas (**Figure S5D**), probably due to the induction of diencephalic/mesencephalic-like domains described above. Interestingly, ventral medial ganglionic eminence (MGE) markers (SHH, LHX8 and NKX2-1) were not highly represented in WNTi-FGF8 organoids, whereas lateral ganglionic eminence (LGE)-enriched markers (EBF1, GSX2 and PBX3) were more expressed than in control samples (**Figures 4C,D** and **S5B**), suggesting that FGF8-treated organoids acquire an LGE ventral identity, but not an MGE-like one. Altogether, these data support an FGF8-mediated effect on the acquisition of a ventral (LGE-like) GABAergic identity at the expense of a dorsal glutamatergic identity in telencephalic organoids.

**Figure 4.**
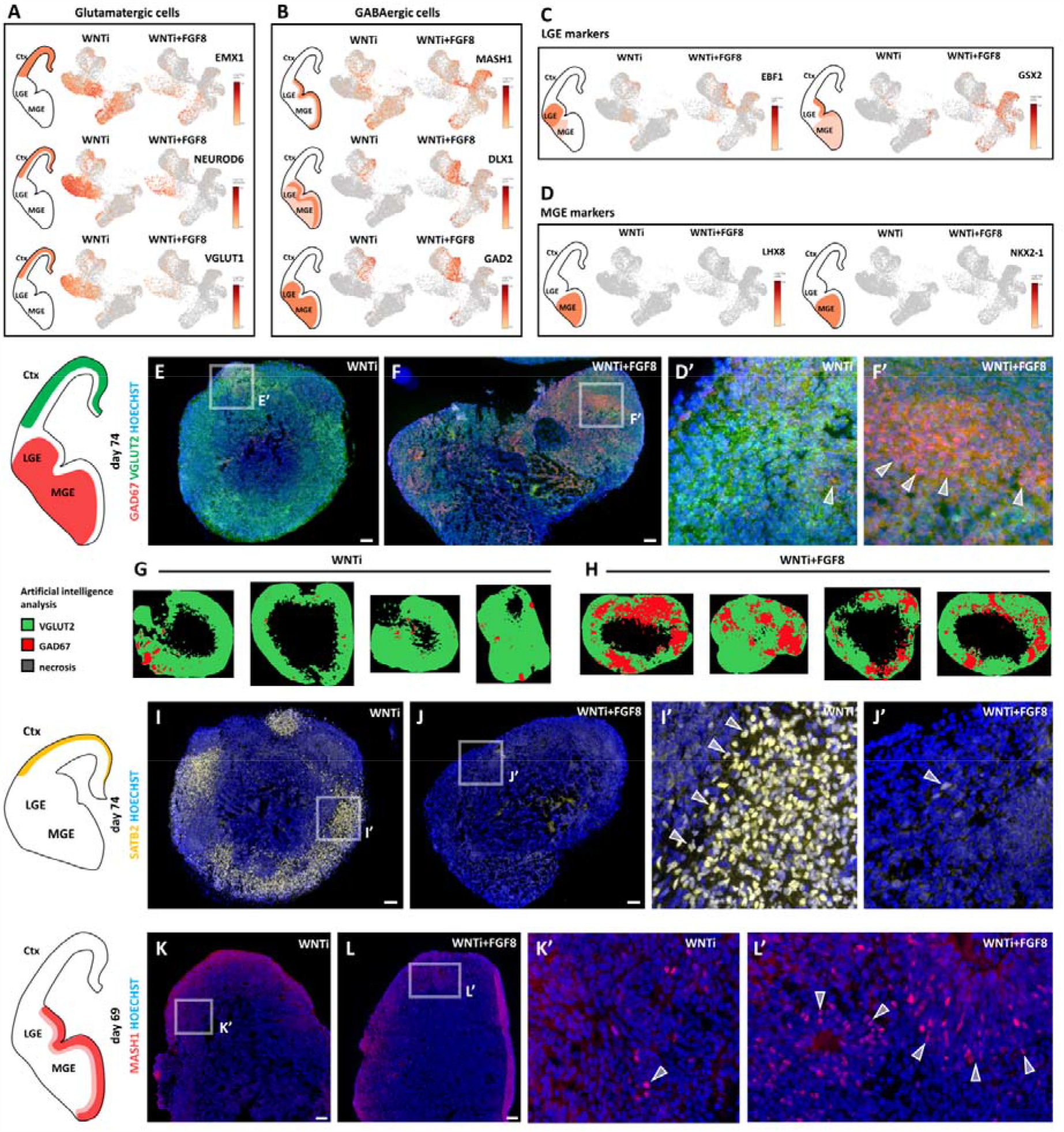
Effects of FGF8 treatment on the on dorso-ventral cellular composition of telencephalic organoids. **A,B)** Expression levels of known markers identifying dorsal glutamatergic NPs and neurons (panel A; EMX1, NEUROD6 and VGLUT1) or ventral GABAergic NPs and neurons (panel B; MASH1, DLX1 and GAD2). **C,D)** Expression levels of known markers identifying ventral GABAergic NPs and neurons of the LGE (Panel C; EBF1 and GSX2) or of the MGE (Panel D; NKX2-1 and LHX8). **E-F’)** GAD67 (red) and VGLUT2 (green) immunostaining in day74 control (WNTi) and treated (WNTi+FGF8) organoids, as indicated. The distribution of these markers in vivo is shown in the brain scheme on the left, while GAD67 pixel intensity quantification is shown in Suppl. Figure 6. **G,H)** Artificial intelligence analysis by HALO software of marker distribution in day74 organoids. Exemplificative images of automatically detected VGLUT2+ (green), GAD67+ (red) and necrotic (black) areas are shown. **I-J’)** SATB2 (yellow) immunostaining in day74 control (WNTi) and treated (WNTi+FGF8) organoids, as indicated. The distribution of SATB2+ neurons in vivo is shown in the brain scheme on the left, while cell density of SATB2+ neurons in these and other samples is quantified in Figure S6. **K-L’)** MASH1 (red) immunostaining in day69 control (WNTi) and treated (WNTi+FGF8) organoids, as indicated. The distribution of MASH1+ ventral progenitors in vivo is shown in the brain scheme on the left, while cell density is shown in Suppl. Figure 6. Scale bars: 100 µm. Abbreviations: Ctx, cortex; MGE, medial ganglionic eminence; LGE, lateral ganglionic eminence.

To directly visualize a change in D/V identity in telencephalic organoids, we performed immunostainings for key GABAergic (GAD67 and MASH1) and glutamatergic (VGLUT2 and SATB2) markers (**Figure 4E-L’**). Consistently with scRNAseq data, FGF8-treated organoids showed increased GAD67 at the expense of VGLUT2 protein expression (**Figures 4E-F’** and **S6A-C**). Artificial intelligence analysis by HALO software estimated a 20% coverage of GAD67+ tissue in FGF8-treated organoids *versus* a 1.8% coverage in control ones (**Figure 4G,H**). Among key dorsal telencephalic genes, double TBR1+ CTIP2+ cells (deep layer cortical neurons) were decreased upon FGF8 treatment (Figure S6D-I), whereas CTIP2+ TBR1-cells were still largely expressed in FGF8-treated organoids. Since these CTIP2+ cells were also expressing GAD67 (**Figure S6J,J’**), we classified them as GABAergic interneurons, similar to what was previously reported in mice^77^. Furthermore, expression of the dorsal cortical marker SATB2 was highly downregulated in WNTi+FGF8 organoids at both transcript and protein levels (**Figures 4I-J’, S5A** and **S6K-M**), in line with a partial loss of dorsal glutamatergic in favor of a ventral GABAergic identity. Among the ventral markers evaluated in scRNAseq, we detected an increased number of MASH1+ ventral progenitors in WNTi+FGF8 organoids compared to WNTi organoids (**Figures 4K-L’** and **S6N-P**). Notably, FGF8-mediated induction of ventral GAD67 and MASH1 markers, and concomitant reduction of dorsal markers VGLUT2, TBR1 and SATB2, were present only in FOXG1+ telencephalic and not in non-telencephalic OTX2+ regions of multi-regional organoids, as supported by comparing sequential cryostat sections of control and treated organoids (**Figure S7A-J’’**). Furthermore, we found that FGF8-mediated downregulation of NR2F1 was present only in FOXG1+ telencephalic regions, whereas NR2F1 was not modulated by FGF8 in non-telencephalic OTX2+ regions corresponding to clusters 8/9 (**Figure S7G-G’’,K,L**). This indicates that FGF8-mediated target gene modulation follows distinct genetic rules depending on the regional identity acquired by NPs and neurons (**Figure S7M**). Altogether, our results point to a strong and telencephalon-specific effect of FGF8 on the D/V (*i*.*e*., glutamatergic *versus* GABAergic) identity of NPs and neurons during human brain development *in vitro*.

Consistently with a change in D/V neuronal composition, four months-old FGF8-treated organoids recorded with a multi-electrode array (MEA) displayed striking differences in terms of spontaneous activity, when compared with non-treated ones (**Figure 5**). Notably, WNTi+FGF8 organoids showed lower spike frequency (firing rate) and amplitude, and a longer recovery time between spikes (interspike interval) compared to WNTi organoids (**Figure 5A,B**). The network analysis (*i*.*e*. the synchronicity of spike events), a read-out of neural network formation, also highlighted differences between the two types of organoids (**Figure 5C,D**). While WNTi organoids showed a consistent degree of network formation, with high number of spikes per burst and high firing rate (**Figure 5C**), WNTi+FGF8-treated ones displayed lower activity levels both in terms of number and frequency of spikes per burst (**Figure 5D**). Nevertheless, synchronous events were still detected in WNTi+FGF8-treated organoids (**Figure 5D**), suggesting a good level of spontaneous circuitry organization despite lower activity levels. Furthermore, axonal tracking identified efficient signal transduction along tracts departing from WNTi organoids and expanding across the MEA electrodes with high conduction velocity (**Figure 5E**). On the contrary, WNTi+FGF8-treated organoids generated normal amplitude signals, however these struggled to travel at high distance (**Figure 5F**). These data indicate that prolonged FGF8 exposure affects neuronal identity (in terms of glutamatergic *versus* GABAergic balance) and function and, consequently, the spontaneous electric activity of neuronal circuits developing in 3D organoids.

**Figure 5.**
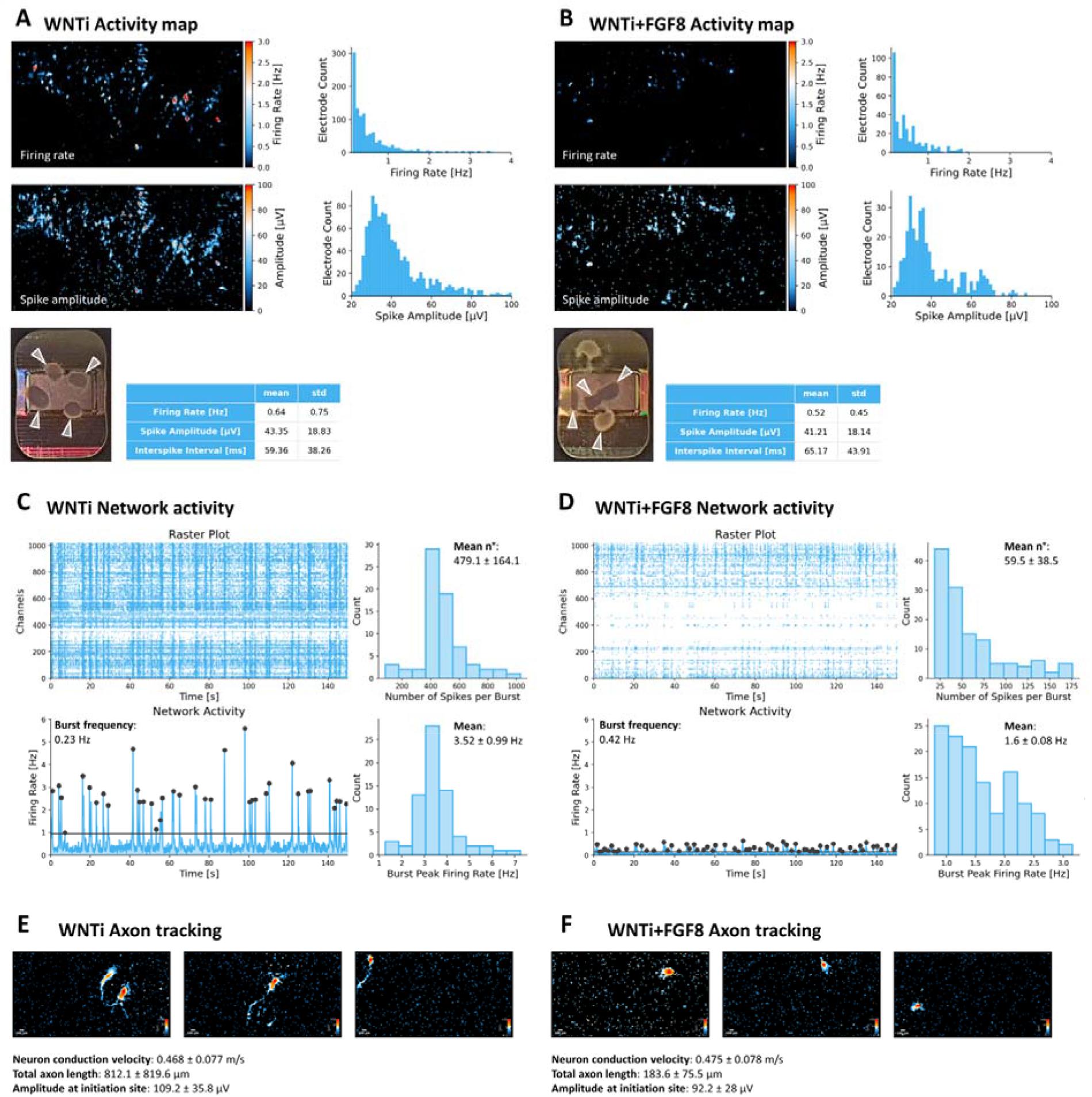
Electrophysiological spontaneous activity of control and FGF8-treated 3D organoid networks. **A,B)** Spontaneous activity maps showing the mean firing rate and the mean spike amplitude of 4 months old control (WNTi; A) and treated organoids (WNTi+FGF8; B), two weeks after plating on high-density MEA chips. Histograms show the firing rates and spike amplitudes of single events captured by electrodes, while mean values are shown in the tables. White arrowheads in electrode images point at organoids adhering totally or partially on the MEA recording surface. **C,D)** Temporal raster plots showing firing events recorded by the 1024 most active electrodes (upper graphs) and their synchronicity suggestive of network activity (lower graphs), as detected from 4 months old control (WNTi; C) and treated organoids (WNTi+FGF8; D), two weeks after plating. Graphs show the number of spike per burst and the burst peak firing rate. **E,F)** Exemplificative images of neuronal tracts extending from organoids on MEA surface, as detected by the automatic axon tracking function, in WNTi (E) and WNTi+FGF8 (F) samples.

### FGF8 alters dorso/ventral specification of glutamatergic and GABAergic populations

As the abundance of cells in specific clusters was highly impacted upon FGF8 treatment, we reasoned that the modulation of specific D/V genes (**Figure 4**) could result from two overlapping factors: (i) a different number of cells per cluster and (ii) a different expression level of D/V genes in a given cellular population. To untangle these two parameters, we performed an analysis of differentially expressed genes (DEGs) on specific cell populations (clusters), by comparing WNTi to WNTi+FGF8 organoids, and visualized the most strongly up- or down-regulated genes in volcano-plots (**Figure 6**). This analysis has the advantage of normalizing the expression level of a gene to avoid any possible bias due to different cell abundance in clusters between control and treated organoids.

**Figure 6.**
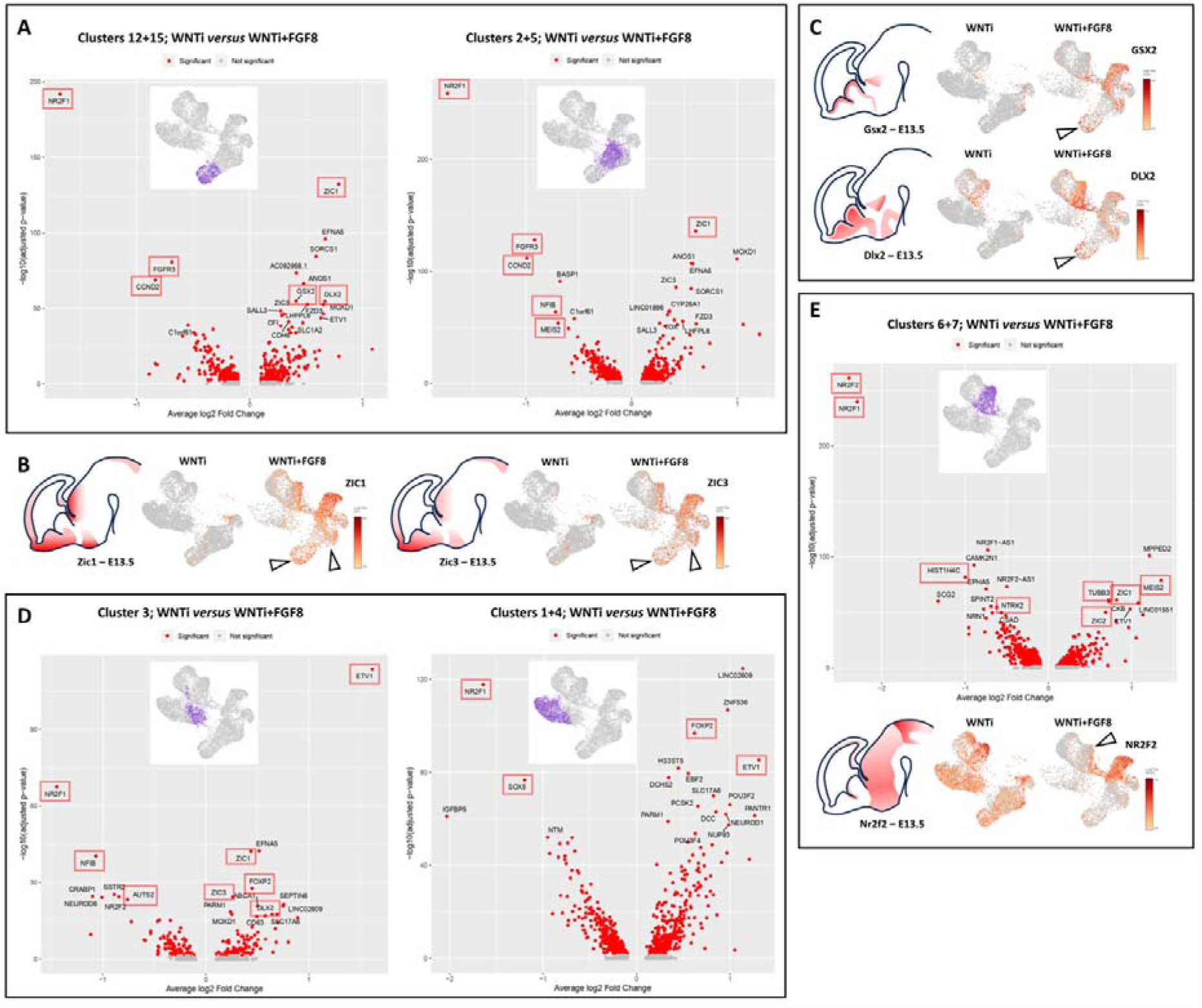
Analysis of differentially expressed genes (DEGs) in glutamatergic and GABAergic progenitors and neurons upon FGF8 treatment. **A)** DEG analysis comparing control (WNTi) and treated (WNTi+FGF8) organoids and highlighting most strongly (x-axis, average log2 fold change) and most significantly (y-axis, adjusted p-value in -log10) differentially expressed genes in clusters 12/15 (proliferating glutamatergic progenitors; left volcano plot) or in clusters 2/5 (non-proliferating glutamatergic progenitors; right volcano plot). Red dots correspond to significantly (adjusted p-value < 0.05) regulated genes, but only the names of the 20-most significantly ones are displayed (See supplementary material for full list of DEGs). **B)** Expression level of ZIC1 (left) and ZIC3 (right) in UMAP projections of WNTi and WNTi+FGF8 samples, as indicated. Black arrowheads point to increased ZIC1 and ZIC3 expression in glutamatergic progenitor clusters upon FGF8 treatment. **C)** Expression level of GSX2 (upper panel) and DLX2 (lower panel) in UMAP projections of WNTi or WNTi+FGF8 samples, as indicated. Black arrowheads point to increased GSX2 and DLX2 expression in proliferating glutamatergic progenitors upon FGF8 treatment. **D)** DGE analysis comparing control (WNTi) and treated (WNTi+FGF8) organoids and highlighting most significantly regulated genes in cluster 3 (early differentiating glutamatergic neurons; left volcano plot) or in clusters 1/4 (differentiated glutamatergic neurons; right volcano plot). **E)** DEG analysis comparing control (WNTi) and treated (WNTi+FGF8) organoids and highlighting most significantly regulated genes in clusters 6/7 (volcano plot), corresponding to GABAergic neurons. The panel below shows the expression level of NR2F2 in UMAP projections of WNTi and WNTi+FGF8 samples, as indicated. Black arrowhead points to decreased NR2F2 expression in GABAergic cells upon FGF8 treatment. Highlighted in red boxes are FGF-target genes or genes referenced in Omim as disease-related genes.

By comparing WNTi and WNTI+FGF8-treated samples on proliferating (clusters 12/15) and non-proliferating (clusters 2/5) glutamatergic progenitors (**Figure 6A**), NR2F1 turned out to be the most strongly down-regulated gene upon FGF8 treatment, both in terms of fold change and p-value. Another down-regulated gene in glutamatergic progenitors was FGFR3, which together with NR2F1 is one of the main targets of FGF8 signaling and one of the key effectors regulating neocortical areal identity^25,78,79^. Enrichment analysis of gene ontology (GO) terms showed that many DEGs belonged to cell proliferation (DNA replication) or differentiation (nervous system development; cell differentiation) categories (**Figure S8**). Among FGF8-induced genes, ZIC1 and ZIC3 (**Figure 6A,B**) are known to maintain neural precursor cells in an undifferentiated state in the mouse medial telencephamon^80^. Most importantly, genes that are normally expressed in ventral progenitors only, such as DLX2 and GSX2, were significantly upregulated in dorsal glutamatergic progenitor clusters (**Figure 6A,C**), suggesting mis-specification of glutamatergic progenitors towards a ventral identity. Post-mitotic differentiating (cluster 3) and differentiated (clusters 1/4) glutamatergic neurons (**Figure 6D**) confirmed a clear imbalance in the expression of D/V genes, *i*.*e*., a reduction of glutamatergic markers (NF1B, NEUROD6 and SOX5) associated with induction of DLX2, validating that FGF8 can affect the establishment of a glutamatergic molecular network by inducing the expression of ectopic GABAergic markers. In general, DEGs in clusters 1/3/4 belonged to neural differentiation GO categories (axonogenesis; neurogenesis; neuron differentiation; axon guidance; cell differentiation, among others) (**Figure S8**), consistently with a post-mitotic neural identity of these clusters. Among FGF8-induced genes in glutamatergic neurons, we found ETV1 (also called ER81)^22^, a target gene considered to be an internal control of FGF8 efficiency, and again NR2F1 as the most significantly down-regulated gene (**Figure 6D**). Notably, among other factors modulated by FGF8 in both NPs and neurons, we found AUTS2, NFIB, ZIC1, ZIC3, FOXP2, CCND2, MEIS2 and SOX5, known to be mutated in NDDs (**Figure 6A,D**). Finally, we focused on DEG changes in ventral GABAergic cells (clusters 6/7) (**Figure 6E**). Gene enrichment analysis revealed GO categories connected with neuronal differentiation and migration (forebrain development; CNS development; neuron migration; nervous system development, among others) (**Figure S8**). Expression levels of NR2F1, NTRX2, TUBB3, ZIC1, ZIC2 and MEIS2, were also affected by FGF8 treatment, and, notably, the NR2F1 homolog NR2F2 also appeared as a highly modulated target gene by FGF8 (**Figure 6E**). Together, our DEG analyses suggest an FGF8-dependent mis-specification of glutamatergic and GABAergic progenitors and neurons, and the concomitant dysregulation of several developmental and NDD-related genes.

### FGF8 treatment partially affects A/P neocortical identity and expression of areal-specific factors

As the DEG analysis showed downregulation of the posterior factors NR2F1 and FGFR3 and upregulation of anterior genes such as ZIC1 and ZIC3 upon FGF8 treatment, we reasoned that FGF8 could modulate A/P identity of telencephalic cells, besides the D/V one. Notably, NR2F1 is located at the top of a hierarchy regulating other regionalized factors such as EMX2, FGFR3, SP8 and PAX6^31^ (**Figure 7A**). VoxHunt Similarity brain maps showed that FGF8-treated organoids still had a high similarity score with the dorsal telencephalon of anterior-most brain sections, while they almost completely lost any similarity to posterior-most areas (**Figure 7B**), indicating that FGF8 treatment mainly sustains the expression of anterior cortical genes. To further explore the effect of FGF8 on A/P patterning, we quantified the percentage of cells positive for selected A/P master genes^83^ (**Figure 7C-E**), in both glutamatergic NP (clusters 2/5/12/14/15) or neurons (clusters 1/3/4). Anterior markers, such as PAX6 and ER81 (ETV1), showed increased expression in progenitors and neurons (for ER81) upon FGF8 treatment (**Figure 7D-F**). Vice versa, and consistent with the upregulation of anterior markers, posterior markers such as NR2F1 and FGFR3 were efficiently down-regulated by FGF8 in both NPs and neurons (**Figure 7D-F**). However, other anterior (ETV5) and posterior genes (CRYM, TSHZ2 and ODZ3) did not show strong changes (**Figure 7D,E**). Unexpectedly, the percentage of cells expressing the posterior gene WNT7B was slightly increased upon FGF8 treatment (**Figure 7D,E**), and the posterior areal patterning gene EMX2 was downregulated in progenitors but upregulated in early differentiating neurons (**Figure 7D,E**). These data indicate a partial effect of FGF8 on the A/P identity of telencephalic organoids, with only a subset of known areal-specific genes being efficiently modulated in glutamatergic clusters. We thus reasoned that a more high-throughput, unbiased analysis was necessary to evaluate the A/P identity of telencephalic organoids. To this aim, we used SingleR to compare the transcriptomic expression profile of control and FGF8-treated organoids against a published reference dataset with distinct human fetal brain areas at GW18 (parietal, motor, pre-frontal, somatosensory, temporal and visual cortices)^84–87^ (**Figure 7G**). Notably, WNTi and WNTi+FGF8 organoid samples (each subdivided into NP clusters or differentiated neuron clusters) showed high similarity to GW18 pre-frontal and somatosensory cortices (**Figure 7G,** left graph). Particularly, by visualizing the SingleR annotation score between organoid samples and fetal areas (**Figure 7G,** right graph), NPs (both control and FGF8-treated) largely resembled the fetal somatosensory cortex, while differentiated neurons were more similar to the pre-frontal or the somatosensory cortex. Despite this, we found a detectable effect of FGF8 on the A/P identity consisting in a slightly higher similarity of WNTi+FGF8 NPs to the pre-frontal transcriptional profile, and a lower similarity of WNTi+FGF8 NPs and neurons to the fetal visual cortex and temporal areas (**Figure 7G,** right graph). Together, our data suggest an effect of FGF8 in areal patterning by inducing anterior (pre-frontal) cortical identity at the expense of a posterior (visual) one.

**Figure 7.**
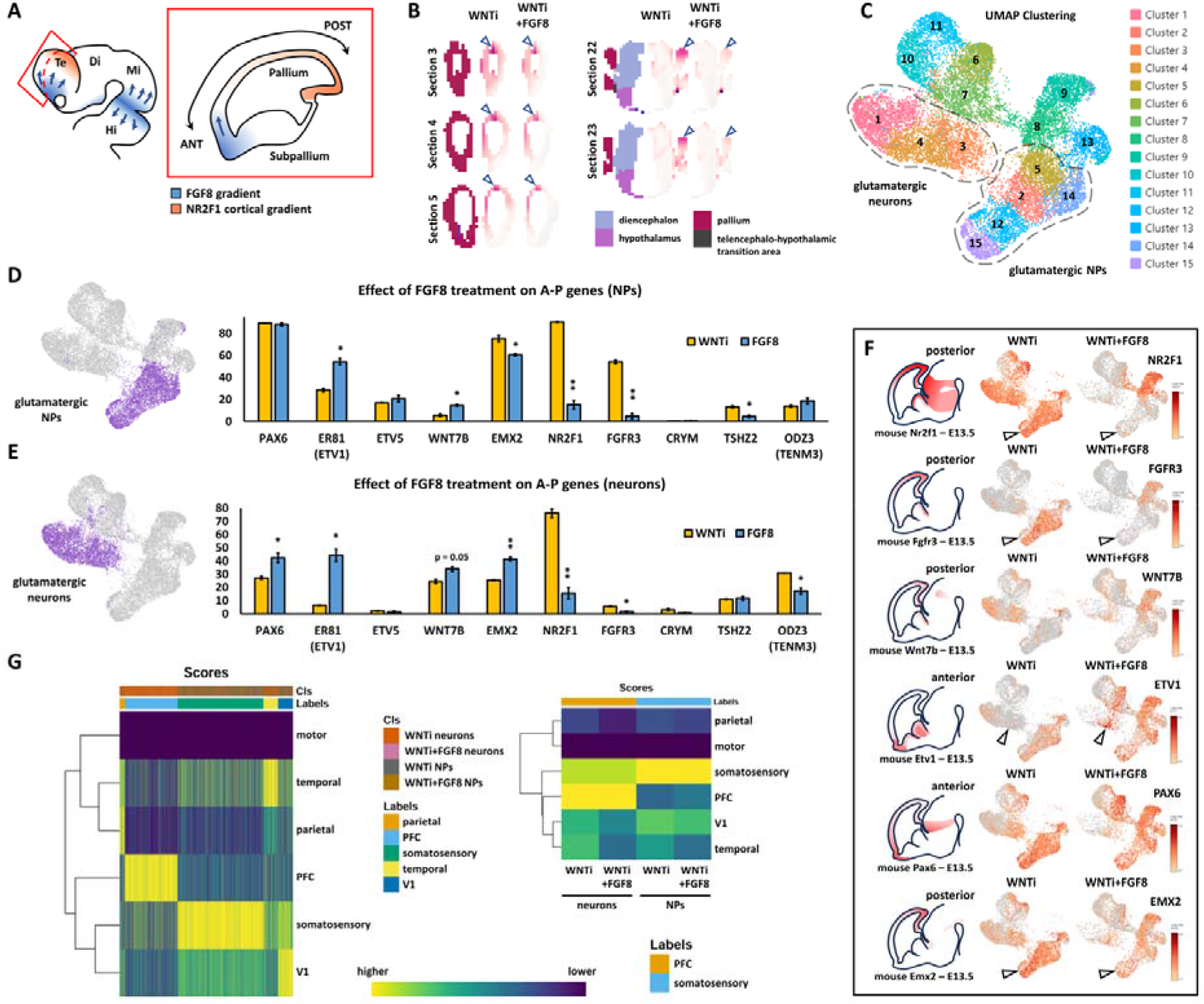
FGF8-dependent acquisition of different antero-posterior areal identities in human telencephalic organoids. **A)** Schematic drawing of the mammalian developing brain showing FGF8 sources (blue) in the anterior telencephalon and at the midbrain/hindbrain border, and their presumable diffusing gradients (arrows). The red inset shows a sagittal section of the telencephalon with the two opposite FGF8 (blue) and NR2F1 (orange) gradients. **B)** VoxHunt Similarity Map showing similarity correlation index (white to violet color code) of control (WNTi) and treated (WNTi+FGF8) organoids on virtual brain coronal sections. The color code in the first left column of brain virtual sections allows to identify different brain regions, listed in the legend at the bottom. Blue arrowheads point to the dorsal-most region of the pallium. **C)** Glutamatergic neural progenitor (NPs; clusters 2/5/12/14/15) and glutamatergic neurons (clusters 1/3/4) are highlighted on the UMAP projection of day69 organoid scRNAseq data. **D,E)** The images on the right show clusters that were selected for the analysis, as described in C. Graphs show the percentages of cells positive for antero-posterior (A-P) cortical markers in the two highlighted cluster groups, NPs (top graph) and neurons (bottom graph). The percentage of positive cells for anterior (PAX6, ER81, ETV5) and for posterior (WNT7B, EMX2, NR2F1, FGFR3, CRYM, TSHZ2 and ODZ3) markers is shown in yellow for control (WNTi) organoids and in blue for FGF8-treated (WNTi+FGF8) organoids. **F)** Expression level of key posterior (NR2F1, FGFR3, WNT7B, EMX2) and anterior (ETV1, PAX6) genes in UMAP projections of WNTi or WNTi+FGF8 day69 organoid samples, as indicated. Black arrowheads in the NR2F1 and FGFR3 UMAP projections point to decreased expression in proliferating glutamatergic progenitors upon FGF8 treatment, while arrowheads in ETV1 UMAP projection point to increased expression in FGF8-treated glutamatergic neurons. **G)** Cell-level (left) and cluster-level (right) heatmaps of the SingleR assignment scores (i.e. the confidence of the predicted labels across the dataset; dark blue to yellow color code) as well as the corresponding inferred annotation for the clusters/cells in the “Labels” top bar. Cell/cluster annotation was obtained by using SingleR to evaluate the similarity between control (WNTi) or FGF8-treated (WNTi+FGF8) samples against a GW18 fetal brain dataset. Glutamatergic progenitors corresponded to clusters 2/5/12/14/15, while neurons corresponded to clusters 1/3/4. The reference dataset corresponds to primary cells dataset published by Speir et al. (2021) and where only cells belonging to GW18 and not to the hippocampus were kept. In the left graph, the “Cls” top bar identify the organoid cells. Note that most of the organoid cells (columns) were annotated as pre-fontal cortex (PFC; light blue) or somatosensory cortex (green), based on transcriptional similarity. In the graph on the right, the average annotation score per sample is shown; control and treated progenitors show high annotation score to somatosensory cortex, while control and treated neurons resemble the PFC. Despite this, note that FGF8 treatment decreased the annotation score to caudal V1 (visual) and temporal areas, while slightly increasing the annotation score to the rostral PFC, indicating anteriorization of the cell identity.

## DISCUSSION

### A hybrid 2D/3D cerebral organoid protocol allows rapid generation of telencephalic progenitors and neurons

In the last decade, 3D human organoids have emerged as a promising tool to model both normal and pathological embryonic brain development. Here, we adapted previous protocols to propose a new strategy combining 2D efficient NP induction^63^ with 3D culture in spinning bioreactors^67^. While dual SMAD inhibition in 2D allows fast neural induction in 7 days only, following steps in 3D lead to the formation of brain-like cellular architecture, notably rosettes or neural epithelia, which gradually develop into neurons organized in a cortical plate-like structure. Since early NPs are dissociated and re-aggregated to create 3D embryoid bodies, we propose that this hybrid 2D/3D method could be of particular interest for experimental approaches aiming at mixing different NP types or cells with distinct genetic backgrounds. It must be noted that the use of a ROCK inhibitor was required during early NP dissociation, as cells died in its absence in the 24 hours following dissociation, indicating that early human progenitor cells are prone to apoptosis when losing cell contacts, in a similar way to undifferentiated hiPSCs. In late culture samples, our scRNAseq data highlighted the presence of distinct types of human NPs, including HOPX+ outer RG cells^71^ and, particularly, CRYAB+ apical truncated RG cells, which to our knowledge, have not been identified in so many other protocols^72^. Such a high cellular heterogeneity could be presumably promoted by the dual Smad inhibition approach coupled with WNT inhibition, a condition shown to enhance NP amplification and diversity^88^.

### FGF8-induced cellular heterogeneity leads to the formation of segregated co-developing domains in multi-regional organoids

Regionalization of the embryonic brain is achieved through a multi-step process that operates sequentially and/or simultaneously, and is mainly controlled by localized sources of various signaling molecules that act as organizing centers by patterning neighboring fields to create molecularly distinct domains^1,2^. From E8.5 in the mouse, FGF8 is expressed at the boundary between the midbrain and the hindbrain, where it regulates posterior brain patterning and strongly induces midbrain identity^10,89,90^. This is why protocols aiming at producing midbrain neurons *in vitro* employ early FGF8 treatment^63,91^. Consistently, we found that the early addition of FGF8 in the culture media abolished FOXG1 expression, *i*.*e*., prevented telencephalic induction. Therefore, we treated organoids after a minimal period of 10 days of neural induction, to avoid early interference with telencephalic development. Despite this, some diencephalic/mesencephalic-like areas still formed in our organoids, which could be due to the presence of unspecified NPs at day10, still able to respond to the mesencephalon-inducing function of FGF8. We propose that FGF8 in human cells can play multiple roles *in vitro*, depending on the competence, developmental state, and regional identity of NPs exposed to it; early in time, when acting on unspecified progenitors, FGF8 is a strong inducer of posteriorized identity, able to specify diencephalic/mesencephalic-like discrete domains in brain organoids; however, later in development, after efficient expression of FOXG1 and consolidation of a telencephalic fate, FGF8 becomes a regulator of A/P and D/V identities in forebrain cells. Interestingly, the ability of FGF8 to modulate target genes, e.g. NR2F1, can also change in function of the regional identity of FGF signaling-exposed cells.

The co-presence of different regional domains in FGF8-treated organoids (notably dorsal and ventral telencephalon, as well as diencephalon/mesencephalon) is very interesting, as one of the current challenges in the field of *in vitro* models of human brain development is to successfully reproduce the interaction between distinct brain regions. Unguided organoids spontaneously develop multiple brain regions^58–60^, but they do so in a stochastic manner, lacking reproducibility from batch to batch^74^. A more reliable strategy consists in using signaling molecules in the form of morphogens and chemical drugs in the culture medium, controlling patterning of the organoids into, for example, cortex, ventral telencephalon, hypothalamus, and thalamus. However, organoids specified to distinct identities require additional manipulation to be fused in multiwells and/or embedded together in matrigel to form “assembloids”^92^, a time-consuming step possibly adding further variability, even if necessary to study neuronal migration or to enable the formation of inter-region neuronal connections. Here, we show that the addition of patterning cues (FGF8 in this case) in the culture medium, even when provided in a non-polarized manner, is sufficient to instruct additional regional fates that co-exist and co-develop in the same organoid while maintaining spatial segregation. In particular, we found FOXG1+ telencephalic areas which contained both TBR1+ dorsal cortical neurons and ventral GABAergic cells, co-existing with more posterior OTX2+ FOXG1-diencephalic/mesencephalic-like domains. Hence, by increasing organoid complexity, FGF8 signaling allows the development of more biologically realistic models of human brain assembly *in vitro*. Multi-regional organoids offer the potential to understand how different parts of the brain self-organize and interact, bypassing the need to manually assemble pre-patterned organoids into assembloids. We propose that the simple addition of instructing cues (morphogens) to culture media could increase the complexity of brain organoids, providing a compromise between maintaining a certain degree of spontaneous self-organization while inducing multiple brain regions in a reproducible manner.

### Control of D/V and A/P telencephalic identity by FGF8 signaling

Our data on human cerebral organoids show that the major effects of FGF8 exposure on organoid NPs and neurons within telencephalic territories, i.e., FOXG1+ cells, were induction of ventral GABAergic at the expense of dorsal glutamatergic cells, and induction of anterior at the expense of posterior neocortical genes, indicating an FGF8-mediated effect on D/V and A/P identities. FGF8 protein has a graded distribution in the telencephalon and acts as a morphogen to drive diverse cellular responses distal to its anterior source, the ANR. In mice, reducing Fgf8 dosage causes progressive telencephalic hypoplasia^17^. Indeed, Fgf8 hypomorphic and conditional mouse mutants have a smaller telencephalon, due to reduced proliferation, high apoptosis and alterations in the expression of areal patterning genes, such as Nr2f1, Pax6, Emx2 and Sp8^17,18,78^. By regulating adjacent signaling centers expressing Bmp4, Wnt8b and Shh, severe hypomorphic and conditional *Fgf8null* mutants have strongly reduced antero-ventral structures, whereas conditional *Fgf8* mutants in which *Fgf8* is inactivated at a slightly later stage, show a milder phenotype but still a reduced frontal cortex, diminished ventral structures and an expanded dorso-posterior molecular profile^17,78^. Thus, severe phenotypic alterations and cross-regulation between anterior (FGF), dorsal (BMP, WNT) and ventral (SHH) patterning centers in genetic loss-of-function (LOF) animal models, mask the unique role of FGF8 signaling in telencephalic development *in vivo*. A system in which FGF8 signaling can be modulated in a controlled manner is therefore preferable. This has been obtained in mouse short-term E9.5 telencephalic explants treated with FGF8-soaked beads, in which expression of some ventral markers such as Mash1 and Dlx2 was induced, while the dorsal Emx1 cortical marker was repressed^93^. Moreover, *in utero* electroporation of an Fgf8-expressing vector can re-specify positional area identity in the neocortex of E11.5 embryos, without affecting cortical growth^19^. Distinct experimental outcomes regarding either A/P or D/V identity suggest that FGF8 acts differently according to the developmental stage and target cellular competency^94^.

In this study, we were able to follow the exclusive and long-term effect of FGF8 signaling in cerebral human organoids by allowing proper differentiation of FOXG1+ telencephalic cells and assessing potential molecular and cellular changes at different stages. We found that FGF8 treatment starting at day10-11 induced FGF-signaling target genes without significantly affecting FOXG1 expression. Hence, we assumed that this was a proper stage for testing the effect of FGF8 in FOXG1+ cells, i.e., soon after having acquired a telencephalic identity. Then, we analyzed by scRNAseq the transcriptomic signature of FGF8-treated *versus* non-treated cerebral organoids and characterized the cellular composition of telencephalic NP and differentiating neurons. The expression of CRYAB indicated that our cerebral organoids might have reached a stage equivalent to an embryonic age of GW16.5 *in vivo* (end of the first trimester). We found a clear FGF8-dependent effect on D/V specification, notably an FGF8-mediated induction of ventral telencephalic genes in FOXG1+ cells, suggesting that one of the first actions of FGF8 signaling might be the regulation of ventral fate during telencephalic development. In accordance with the induction of ventral genes, such as MASH1, DLX2, PBX3, and the concomitant reduction of dorsal ones (EMX1, NEUROG2, SOX5, LHX2), FGF8-treated organoids showed an imbalance between glutamatergic neurons expressing NEUROD6, NEUROD2, CTIP2, TBR1, SATB2, NF1A and GLUT1, and GABAergic and/or striatal neurons expressing EBF1, GSX2, PBX3 and GAD1. Accordingly, our functional assay on MEA showed decreased spontaneous network activity in FGF8-treated organoids and decreased electric signal propagation along FGF8-treated axons when compared to non-treated ones. A similar electrophysiological profile, i.e., lower firing rate, network burst rate and signal conduction, is compatible with a reduction in activity due to a decrease in excitatory glutamatergic neurons and an increase in inhibitory GABAergic neurons *in vitro*^*95*^. Strikingly, we found that some of these dysregulated genes are specifically expressed in the LGE, a region localized just ventral to the neocortex, while MGE markers, such as SHH, LHX8, and NKX2-1, were not affected upon FGF8 treatment. These data strongly support an FGF8-mediated effect on the acquisition of a ventral (LGE-like) GABAergic identity at the expense of a dorsal glutamatergic one in telencephalic organoids. They also strongly indicate that FGF8 signaling at this stage of human development is involved in specifying LGE-but not MGE-derived cells. This is an unexpected finding that, to our knowledge, has never been described before in animal models and points to either a human-specific trait of FGF8 signaling or to the fact that Fgf8 genetic LOF models induce such a severe loss of ventral structures that the specific effect on the LGE or MGE domains cannot be clearly sorted.

FGF8 is also known as a key regulator of anterior *versus* posterior identity, leading to the specification of different cortical areas in the mouse^96^. FGF8 gradient diffuses from anterior to posterior along the neocortical epithelium and specifies frontal while inhibiting more occipital cortical areas, such as the visual cortex. Although human cerebral organoids do not form segregated functional areas as is the case in the developing neocortex^83,84^, we found that FGF8 can down-regulate the expression of posterior cortical genes in human organoids, such as FGFR3 and NR2F1, while inducing anterior ones, such as ETV1. This indicates that FGF8, besides controlling the D/V identity of neural cells, can also modulate the expression level of key A/P areal-specific factors in human cerebral organoids. However, the global identity of both treated and untreated organoids, based on the similarity of their transcriptional profile to that of human fetal brain areas, remained largely like the somatosensory and pre-frontal human cortical regions, indicating a limited effect on A/P area specification. An alternative strategy or a refinement in terms of FGF8 doses and/or temporal windows could help in more efficiently controlling the areal identity of telencephalic organoids. One possible way to achieve this could be the use of a polarized source to reproduce a more physiological FGF8 gradient in organoids. This has been done for SHH signalling^97^, showing that an endogenous signaling center can induce distinct D/V identities in a dose-dependent manner. We would expect that a discrete FGF8 source inside a developing forebrain organoid could expose telencephalic cells to different dosages of diffusing molecules, hence modelling *in vitro* the process of areal patterning more efficiently and more physiologically, by inducing distinct areal identities and eventually neocortical axes in organoids.

In summary, we showed that FGF8 impacts the establishment of D/V and A/P regional identities in telencephalic organoids. Even if we analyzed the effects on the A/P and on the D/V axis separately, it is reasonable to think that A/P and D/V inductions are two closely interconnected processes. Considering that the ANR is placed medially in an anterior and ventral position in the early telencephalon, we could define FGF8 as an “antero-ventral” inducer. This could explain the double effect that FGF8 treatment produces in our cerebral organoids, i.e., an inducer of both ventral (LGE-like) and anterior brain identity.

### FGF8 signaling impacts NDD-related developmental trajectories

By employing long-term treatment of human organoids, we found a correlation between FGF signaling activation and the modulation of several developmental and/or NDD-related genes. Among FGF8-regulated genes, NR2F1 was the most strongly and significantly regulated one, suggesting that it might be one of the major effectors of FGF8 signaling during telencephalic development. Besides NR2F1, other FGF8 responding genes that we detected in the DEG analysis, are also implicated in human brain malformations and/or in NDDs. For example, FGFR3 dysregulation leads to Thanatophoric dysplasia, a fatal form of chondrodysplastic dwarfism, in which the cerebral cortex displays temporal lobe enlargement with abnormal sulci and hippocampal dysplasia, leading to cognitive impairments, such as severe memory deficits and low synaptic plasticity in patients and mouse models^98^. ZIC1, significantly up-regulated in FGF8-treated organoids, is instead implicated in complex syndromes with cortical, callosal and cerebellar malformations associated with impaired intellectual disability^99,100^. Haploinsufficiency of the *NFIB* gene results in macrocephaly and impaired intellectual development, similar to what has been described in mouse *Nfib* mutant mice^101^. Additionally, among FGF8-modulated genes detected by our analysis, we found other disease genes related with autism (AUTS2)^102^, with speech and language disorders (FOXP2)^103^ and with developmental delay or intellectual disability (SOX5)^104^. Finally, FOXG1, normally induced by FGF8, has been implicated in a wide spectrum of congenital brain disorders, including the congenital variant of Rett syndrome, infantile spasms, microcephaly, autism ASD and schizophrenia^105^. However, and probably due to early treatment with FGF8, FOXG1 levels tended to decrease instead of increasing in our organoid model, suggesting that FGF8-mediated positive or negative regulation of FOXG1 in NPs and neurons could be time- and regional-dependent. In summary, we found an FGF8-dependent effect on regional identity with concomitant modulation of several NDD-related gene targets, among which an evolutionary conserved FGF8-NR2F1 molecular axis.

All in all, we propose that FGF8-mediated modulation of key developmental genes steers the developmental trajectories of human brain cells along the D/V and A/P telencephalic axes. Different genes and signaling pathways can converge to orchestrate common cellular and molecular processes, leading to similar or overlapping phenotypes in neurodevelopmental pathologies^106^. We speculate that an FGF8-dependent molecular machinery could, upon alteration, trigger NDD phenotypes by dysregulating the expression of NDD-related genes and/or by converging on basic mechanisms of A/P and D/V neuronal specification during early brain development.

## STAR METHODS

### hiPSCs culture

Human induced pluripotent stem cells (hiPSCs) used in this study are the HMGU1 cell line, kind gift of Dr. Drukker. MTA approval was obtained from the Helmholtz Zentrum München (HMGU), Germany. HMGU1 hiPSCs were cultured on Matrigel-coated plates (Corning, 354234; 5-10 µl matrigel dissolved in 1ml cold DMEM-F12 for each well of a 6-well culture plate) in mTeSR1 medium (STEMCELL Technologies; #85850). Medium was changed daily. HMGU1 cells were passaged with Versene (ThermoFisher scientific; 15040066) as previously described^107^; briefly, cells were washed with 1ml PBS1x (ThermoFisher scientific 14190169; or Sigma D8537), treated with Versene for up to 5 minutes, then detached by gently pipetting with 1ml mTeSR1 medium. Alternatively, when single cell dissociation and cell counting were needed, HMGU1 cells were passaged with Accutase (Sigma; A6964). Single cells passaging required addition of 10µM dihydrochloride ROCK inhibitor Y-27632 (MedChemExpress HY-10583; or Stem Cell Technologies #72304) to prevent apoptosis. ROCK inhibitor was removed 24 hours after dissociation. For immunostaining, cells were grown on glass coverslips (Epredia X1000 Round Coverslip dia. 13mm, Product Code: 10513234) with the same medium and coating treatment.

### hiPSCs quality controls

HMGU1 cells were checked for mycoplasma contamination once per year, while amplifying and preparing multiple cryovial aliquots that were used during the following months. Genetic/chromosomic analysis was performed by using the hPSC genetic Analysis kit (Stem Cell Technologies; #07550) by following the manufacturer’s instructions. Two different hiPSC lines (WT-T12, kind gift of Dr. Magdalena Laugsch, and PGP1, purchased from Synthego) were used as controls to be compared with HMGU1 cells. Briefly, hiPSC cell pellets were lysed with lysis buffer (1M TRIS HCL pH8, 5M NaCL, 0.5M EDTA, 10% SDS in milli-Q H^2^O) supplemented with 10mg/mL Proteinase K (Sigma, 0311580); lysis incubation was performed at 58°C for 30 min. Genomic DNA was obtained by Isopropanol precipitation and 70% Ethanol cleaning. We detected frequent mutations in the HMGU1 cell line, and found that extended culture of hiPSCs caused chromosomal abnormalities to further accumulate in sensitive regions. To prevent this, we did not use HMGU1 cells after passage 26. Finally, pluripotency was checked by OCT4 immunostaining.

### Neural differentiation into telencephalic organoids

HMGU1 cells were seeded in Matrigel-coated 24 well-plates at a density of 25.000-50.000 cells per well and cultured in mTeSR1 medium till high confluency, which is key for efficient neural induction in this protocol^63^. When highly confluent (90-95% surface covered), hiPSCs were washed with pre-warmed DMEM/F12 (ThermoFisher scientific; #31331028) then medium was switched to neural progenitor patterning medium (NPPM). NPPM consisted of DMEM/F12 supplemented with N2 (N-2 Supplement 100X; ThermoFisher scientific 17502-048), B27 (B-27™ Supplement 50X, minus vitamin A; ThermoFisher scientific 12587-010), GlutaMAX (ThermoFisher scientific 35050038), Non-Essential Amino-Acid (NEAA) (ThermoFisher scientific 11140-035), Sodium Pyruvate (NaPyr) (ThermoFisher scientific #11360070), 50µM 2-mercaptoethanol (ThermoFisher scientific 31350-010), 2µg/mL Heparin (Sigma H3149) and Penicillin/Streptomycin (Biozol ECL-EC3001D). The latter could be substituted by Antibiotic/Antimycotic solution (Sigma A5955). Neural induction was boosted by adding a BMP inhibitor (0.25µM LDN-193189; Sigma SML0559-5MG) and a TGFβ inhibitor (5µM SB-431542; Sigma S4317-5MG) for the control condition (“CTRL” in figures); telencephalic induction also required use of a WNT inhibitor (2µM XAV-939; Stem cell technologies 72674) (“WNTi” in figures). On day 7 (max on day 8) of the protocol, cells were dissociated by Accutase (Sigma; A6964) and seeded in 96 U-bottom well plates (Corning #7007; or Sarstedt 83.3925500) at a concentration of 50.000 cells/well, in NPPM medium containing ROCK inhibitor. A gentle centrifugation of the 96-well plate (1 minute at 1.000 rpm) allowed quick accumulation of the cells at the bottom of the wells. The day after, the embryoid bodies (EBs) were collected and transferred in Matrigel droplets (Corning 354234; 70µl Matrigel for 16 EBs) to generate “cookies”, distributed on parafilm and incubated 30 minutes at 37°C for Matrigel to solidify, then transferred in SpinΩ mini-rotors^67^ in NPPM medium (still supplemented with SB+LDN±XAV; 2.5ml per well in a 12-well plate). On day9 (max on day10), most of the medium was removed and replaced with NPPM without SB-431542 and LDN-193189, but still containing WNT-inhibitor. Starting on day10, FGF8 (100ng/ML; Recombinant Human/Mouse FGF-8b Protein; R&D 423-F8-025/CF) could be added to NPPM medium, that was then changed every 2-3 days, to create the treated condition (referred to in the text and figures as “WNTi+FGF8”). If medium showed excessively yellow color indicating pH acidification, cookies were diluted by redistributing in multiple wells to have a maximum of 5 cookies per well. At day 20 medium was switched for Neural Differentiation Medium (NDM) containing 50% DMEM-F-12 medium and 50% Neurobasal medium (ThermoFisher scientific 21103049) supplemented with N2, Glutamax, NEAA, NaPyr, 50µM Beta-mercaptoethanol, 2µg/mL Heparin and Insulin (25 µl per 100ML medium; Sigma #19278). At day30 (max at day35), NDM was substituted with Long-term pro-Survival Medium (LTSM) consisting in Neurobasal medium supplemented with N2, B27 without Vitamin A, Glutamax, NEAA, NaPyr, 50µM Beta-mercaptoethanol, 2µg/mL Heparin, 1% Serum (Fetal Bovine Serum, ThermoFisher scientific 10270-106; inactivated for 30 minutes at 56°C then aliquoted and stored at -20°C), 10ng/mL BDNF (PeproTech #450-02) and dissolved Matrigel (final concentration in the medium: 0.1%). Medium was changed every 3-4 days, until the desired differentiation stage. Starting at day50-60, bigger organoids were sliced with a sterile blade under the stereomicroscope (typically once per month), which allows better survival and reduces the formation of an inner necrotic core^62^. Organoids were cultured up to 110 days.

### Real time RT-PCR

Cell pellets or organoids were spinned in 1.5ml Eppendorf tubes and frozen at -80°C. Total RNA was extracted with NucleoSpin RNA II columns (Macherey-Nagel; 740902.50). RNA quantity and quality were assessed with Nanodrop and gel electrophoresis. For each sample, 200/500ng of total RNA were reverse transcribed using random nonamers (Reverse Transcriptase Core Kit, Eurogentec, RT-RTCK-03); RT-PCR was performed using GoTaq SYBR Green qPCR Mix (Promega A6001) or KAPA SYBR^R^ FAST (KAPA Biosystems; KK4610) on LightCycler (Roche). cDNA (stored at -20°C) was diluted so that each reaction contained 2ng. Amplification take-off values were evaluated using the built-in LightCycler relative quantification analysis function, and relative expression was calculated with the 2^-ΔΔCt^ method as previously described^108^, normalizing to the housekeeping gene GAPDH or β-ACTIN. Standard errors for error bars were obtained from the error propagation formula^108^. For each genotype/time point, 2-3 organoids/cell pellets (biological replicates) were pooled together before RNA extraction, while at least 2 reactions were assembled per sample/gene analyzed during RT-PCR amplification (technical replicates).

**Table 1.**
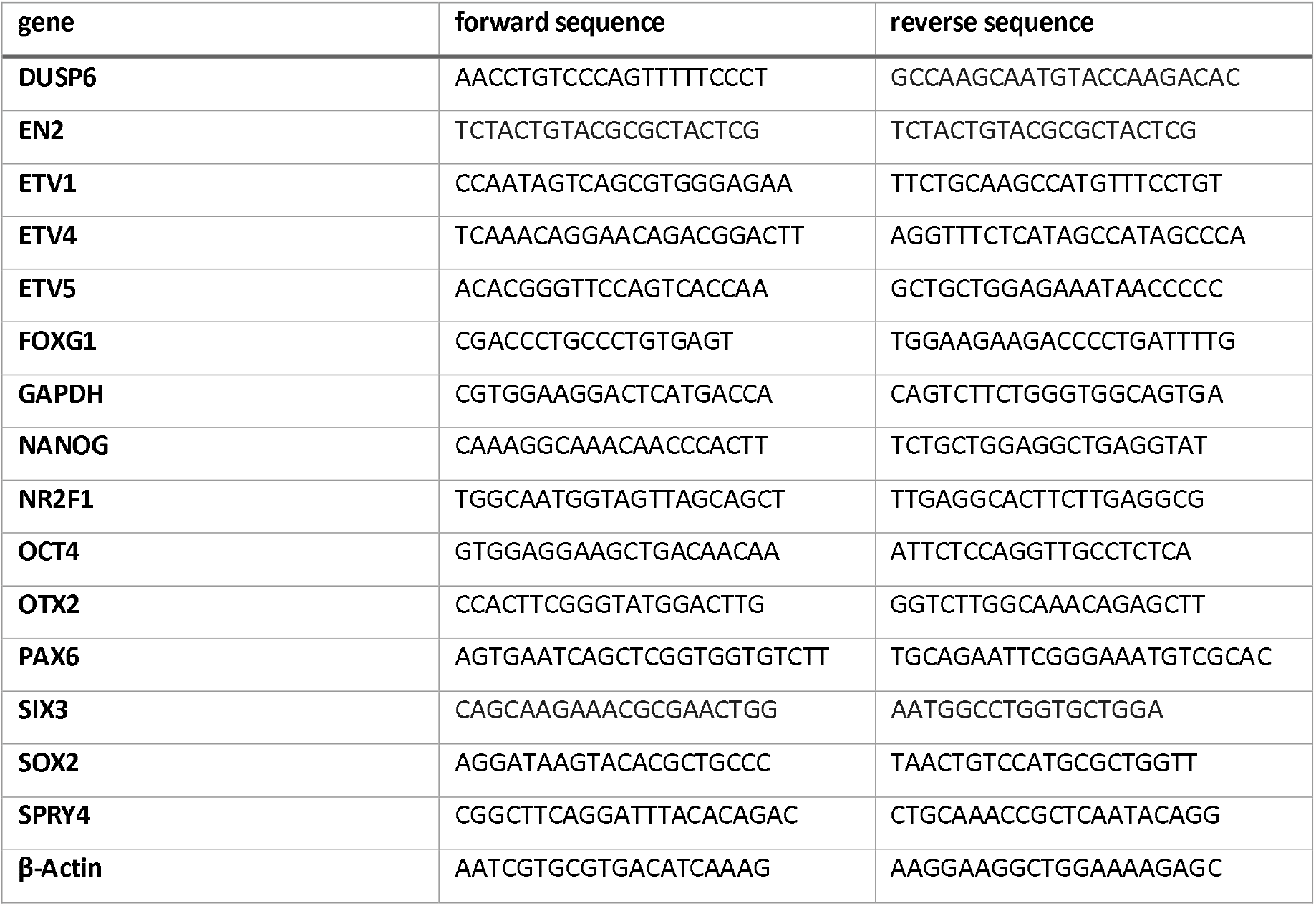
Primer sequences used:

### Immunostaining on cryostat sections

Organoids were collected in 2ml Eppendorf tubes and fixed in 2ml of 4% PFA at 4°C for 3-4 h in gentle agitation, then washed twice with PBS 1X and dehydrated in 10% sucrose (Sigma-Aldrich, S9378-1KG; dissolved in PBS1x; minimum 4 hours to maximum over-night) followed by 25% sucrose (dissolved in PBS1x; overnight at 4°C). Most of the sucrose was removed and substituted with OCT resin (Leica Tissue Freezing Medium, 14020108926; or Cryomatrix ™, 6769006) with a 10 minutes wash with gentle rotation, then organoids were transferred in embedding molds in clean OCT and stored at -80°C. Cryostat 12 µm sections (Leica cryostat, model: CM3050S) were collected on glass slides (ThermoFisher scientific, Superfrost Plus, J1800AMNZ, or VWR SuperFrost™ Plus, 631-0108) and stored at -80°C. Prior starting immunostaining, sections were dried at RT for 10-15 minutes, then washed 2 times with PBS1x (10 minutes each) to remove traces of OCT resin. All antibodies required antigen retrieval prior to incubation (unmasking: 10 minutes at 95°C in pH=6 0.1M sodium citrate solution). After a 5-minute PBS1x wash to remove unmasking solution, pre-blocking solution was added on the slides for 1 hour, containing PBS1x with 5% serum (Sheep serum, Sigma, S2263-100ML; in case of primary antibodies raised in sheep or goat: Newborn Calf Serum, ThermoFisher scientific, 16010-167) and 0.3% Triton (Sigma-Aldrich, T8787-250ML). For primary antibody incubation (from a minimum of 4 hours at RT to a maximum of over-night at 4°C), blocking solution consisted in PBS1x supplemented with 1% serum and 0.1% Triton. Primary antibodies used in this study are listed in Table 2. Alexa Fluor 488, 555, 594, and 647 anti-mouse, anti-rabbit, anti-rat, anti-goat, anti-guinea pig or anti-sheep IgG conjugates (ThermoFisher scientific, all diluted at 1:500) were used as secondary antibodies (incubation time: from a minimum of 2 hours at RT to a maximum of over-night at 4°C). Secondary antibody solution also contained 1:1.000 Hoechst 33342 for nuclei staining (Invitrogen, H3570). After final washes (3 times, 10 minutes each, in PBS 1x), organoid sections were covered with mounting solution (80% glycerol, 2% N-propyl gallate in PBS1x) and glass coverslips, which were sealed with nail polish on the edges. Stained sections were stored at -20°C. Images were acquired at an Apotome Zeiss, using the AxioVision software, and exported as TIF files.

**Table 2.** Primary antibodies used:

| Primary antibody | species | Concentration used | Brand | ref |
| --- | --- | --- | --- | --- |
| CTIP2 | Rat | 1/1000 | Abcam | ab18465 |
| FOXG1 | Rabbit | 1/1000 | Abcam | ab18259 |
| GAD-67 | Mouse | 1/1000 | Millipore | MAB5406 |
| OTX2 | Goat | 1/1000 | R&D | AF1979 |
| MAP2 | Mouse | 1/1000 | Sigma | M4403 |
| MASH1 | Mouse | 1/1000 | BD Biosciences | 556604 |
| **NR2F1** | Rabbit | 1/1000 | Abcam | ab181137 |
| **OCT-3/4** | Mouse | 1/500 | Santa Cruz | sc-5279 |
| **PAX6** | Rabbit | 1/1000 | Millipore | AB2237 |
| **SATB2** | Mouse | 1/1000 | Abcam | ab51502 |
| **SOX2** | Mouse | 1/1000 | R&D | MAB2018 |
| **TBR1** | Rabbit | 1/1000 | Abcam | ab31940 |
| **TUJ1** | Rabbit | 1/1000 | BioLegend | 802001 |
| **VGLUT2** | Guinea Pig | 1/1000 | Millipore | AB2251 |

### Immunostaining of cultured cells

For immunostaining, cells were cultured on Matrigel coated round glass coverslip. The immunostaining protocol is the same used for cryostat sections, except: fixation with 4% paraformaldehyde lasted only 15 min at room temperature; no antigen retrieval was performed; PBS1x for washes always contained 1% serum to limit cell detachment.

### Statistical analysis

All data were statistically analyzed and graphically represented using Microsoft Office Excel software or GraphPad Prism (Version 7.00). Quantitative data are shown as the mean ± SEM. For cell percentage/number quantification after immunostaining, measurements were performed on at least 5 sections coming from 2 to 3 different organoids, unless otherwise stated. Organoid sections with damaged histology were excluded from any further analysis/processing; the inner necrotic core of the organoid, when present, was excluded from counting. Microscope images were processed with Photoshop or ImageJ software, by randomly overlapping fixed-width (100 µm) square boxes on the area of interest [*e*.*g*. organoid surface], then quantifying positive cells inside the boxes. When calculating percentages over the total cell number, the latter was quantified by counting Hoechst+ nuclei, unless otherwise specified. Data were analyzed using the Mann–Whitney U-test or two-tailed Student’s t-test (when comparing two data groups), or by two-way analysis of variance (ANOVA) for comparison of three or more groups. Statistical significance was set as follows: *P<0.05; **P<0.01; ***P<0.001.

### MEA recordings

For MEA recordings, we used the high-density MEA system (HD-MEA) from MaxWell (Model: MaxOne; Chips: MaxOne single-well chips). Prior recordings, organoids were plated on electrodes by following the MaxWell “Brain organoid plating protocol, version 1”, with few modifications. Briefly, MEA chips were cleaned with 1% Terg-a-zyme solution and sterilized with 70% Ethanol. A first coating with Poly-L-ornithine hydrobromide (20µg/ml in distilled water for 5h at 37°C, Sigma P3655) was followed by a second one with Laminin (25µg/ml in PBS1x, Santa Cruz, sc-29012). After a pre-incubation with LTSM medium (prepared as detailed in *Neural differentiation into telencephalic organoids*) to pre-wash the chip surface, organoids were transferred on MEA chips (n=5 organoids per chip) and let deposit by gravity in the incubator, while avoiding any vibration for the next 24 hours. Medium was delicately changed every 3 days during a 2-week incubation on chip; during the second week, medium was gradually switched to LTSM that was prepared using BrainPhys (Stem Cell Technologies, #05790) as a base medium where to dilute the supplements. Last medium change was one day before the recording. Recordings were performed with the MaxOne software, by using in-built protocol “Activity Scan Assay” to detect active areas and to measure mean firing rate and mean spike amplitude, followed by the “Network Assay” to evaluate firing synchronicity and by the “Axon Tracking Assay” to detect signal propagation along axonal tracts. Average data were obtained by recording from the whole chip surface, hence pooling active electrodes from 3/4 organoids per condition.

### Cell dissociation for single cell RNA sequencing and flow cytometry analysis

Day69 control (WNTi) and treated (WNTi+FGF8) organoids were dissociated using an enzymatic Papain Dissociation System (Serlabo technologies LK003150 or Worthington, #LK003150), by following manufacturer’s instructions. Low-bind Eppendorf tubes were used to limit material loss. Two to three organoids per batch were isolated in Eppendorf tubes containing 1 ml of pre-warmed Papain solution supplemented with DNase. Total incubation lasted 70 minutes, with manipulations every ten minutes that consisted in gentle mixing by tube inversion (first 2 rounds) then gentle pipetting (following rounds), ending up with addition of Ovomucoid solution for Papain inactivation. Samples were filtered (Cell Strainer 40µm Nylon, FALCON, 352340) to remove undissociated cells, spinned down (1.200 rpm, corresponding to 170 *g*, for 5 minutes) and resuspended in PBS1x supplemented with 0.1% BSA (Jackson, 001-000-161). Cell viability upon dissociation was checked by Propidium Iodide (PI) staining in flow cytometry in an independent experiment, where Papain was compared with Accutase and with ReLeSR (Stem Cell Technologies, #100-0483). For flow cytometry, cells were stained with Propidium Iodide (40 µg/ML, Sigma, P4170) in PBS1x for 15 minutes at RT, washed twice with PBS1x supplemented with 1% Serum, then analysed with BD LSRFortessa and FACSDiva software (Becton Dickinson) to measure dying cells which incorporated PI. Cells were analysed on the basis of 10.000 total events (debris excluded). Cell viability was tested again by Trypan blue staining just prior cell counting for library preparation, and samples were considered suitable for single cell RNA sequencing only when damaged cells were ≤10% of the total population.

### Single cell RNA sequencing: library preparation and sequencing

Single cell RNA sequencing experiment was performed by using 10X Genomics technology. Dissociated cells were processed following the manufactures’ instruction of Chromium Next GEM Single Cell 31 Reagent Kits v3.1 (Dual Index). Briefly, they were resuspended in ice-cold PBS containing 0.1% BSA at a concentration of 1.000 cells/μl, and approximately 17,400 cells per channel (corresponding to an estimated recovery of 10,000 cells per channel) were loaded onto a Chromium Single Cell 31 Chip (10x Genomics, PN 2000177) and processed through the Chromium controller to generate single-cell gel beads in emulsion (GEMs). scRNA-seq libraries were prepared with the Chromium Single Cell 31 Library & Gel Bead Kit v.2 (10x Genomics, PN-120237). Final cDNA libraries were checked for quality and quantified using 2100 Bioanalyzer (High Sensitivity DNA Assay; Agilent Technologies). Libraries were sequenced using Illumina HiSeq 4000 system in Paired-End mode with 100 or 28 bases for read 1 and 100 bases for read 2, following Illumina’s instructions. Image analysis and base calling were performed using RTA version 2.7.2 and Cell Ranger version 3.0.2.

### Single cell RNA sequencing: analysis Primary analysis

FastQ files of each sample were processed with Cell Ranger count pipeline (10X Genomics) version 6.1.1, that performs alignment, filtering, barcode counting, and UMI counting, using the pre-processed Homo sapiens reference GRCh38-2020-A (GENCODE v32/Ensembl 98) from 10X Genomics. Data were then aggregated using Cell Ranger aggr tool which normalizes counts to the same sequencing depth and then recomputing the feature-barcode matrices and analysis (dimensionality reduction, *i*.*e*. UMAP, and clustering) on the combined data (24,651 cells).A subset of cells expressing endoderm or mesoderm markers (1,662 cells, corresponding to 6.7% of the total population) and a subset of cells expressing high levels of stress markers (4,399 cells, corresponding to 17.8% of the total population) were excluded, and analysis (dimensionality reduction and clustering) was performed again after exclusion of these non-neural and/or sub-optimally differentiated cell populations using Cell Ranger reanalyse tool, resulting a set of 18,590 cells regrouped in 15 clusters. More information about Cell ranger software can be found on the manufacturer website (https://support.10xgenomics.com/single-cell-gene-expression/software/overview/welcome).

### Trajectory analysis

Count data were normalized with the log normalize method using Seurat [REF] R package version 4.0.5 then filtered by keeping only genes with more than 5 UMI counts detected in at least 10 cells. Four analyses were performed: one with all cells (clusters 1 to 15), final data used in the analysis contained 18,590 cells and 7,133 genes; one for ventral progenitors (only WNTi+FGF8 cells belonging to clusters 6, 7, 8, 9 and 13) with 4,608 cells and 5,382 genes; one for dorsal progenitors (only WNTi cells belonging to clusters 1, 2, 3, 4, 5, 12, 13, 14 and 15) with 8,246 cells and 5,827 genes; and the last one for all progenitors (dorsal + ventral; cells belonging to clusters 2, 5, 8, 9, 12, 13, 14 and 15) with 8,876 cells and 6,452 genes. Trajectory inference analyses were performed using slingshot method^109^ implemented in the dyno package^110^.

### VoxHunt analysis

Similarity maps between single-cell data and the public Allen Developing Mouse Brain Atlas expression data, stage E13, were computed using the VoxHunt R package version 1.0.1^73^.

### Analyses of differentially expressed genes (DEGs) and Gene enrichment

Count data were normalized with the log normalize method using Seurat R package version 4.0.5 before performing the analysis^111^. Differential gene expression analyses, realized with Seurat, between clusters of interest are performed using a Wilcoxon test (whose performance for single-cell differential expression analysis has been evaluated in a previous report^112^) and resulting p-values were adjusted for multiple testing using a Bonferroni correction. Enrichment analyses were performed on differentially expressed genes previously identified using cluster Profiler R package version 4.2.0 with Gene Set Enrichment Analysis (GSEA) method^113^. Genes were ranked by their log2 Fold-Change. Enrichment analyses are performed on the three domains of GO (Gene Ontology) terms: biological process; molecular function; and cellular component.

### Cluster annotation with SingleR

Annotation of Neurones_CTRL (clusters 1, 3 and 4 of WNTi cells), Neurones_FGF8 (clusters 1, 3 and 4 of WNTi+FGF8 cells), Progeniteurs_CTRL (clusters 2, 5, 12, 14 and 15 of WNTi cells) and Progeniteurs_FGF8 (clusters 2, 5, 12, 14 and 15 of WNTi+FGF8 cells) cells at the cluster and cell levels were performed with SingleR version 2.0.0 R package^114^ using a previously published primary cell dataset^87^. In this reference dataset, only cells corresponding to GW18 age and not to hippocampus area were kept. Before performing the annotation both datasets were normalized using the LogNormalize method implemented in the Seurat R package version 4.3.0^115^. *Supplementary Figure 11E* shows a heatmap of the SingleR assignment scores as well as the corresponding inferred annotation for the clusters/cells. Scores allows users to inspect the confidence of the predicted labels across the dataset. Ideally, each cell/cluster (*i*.*e*., column of the heatmap) should have one score that is obviously larger than the rest, indicating that it is unambiguously assigned to a single label. A spread of similar scores for a given cell/cluster indicates that the assignment is uncertain, though this may be acceptable if the uncertainty is distributed across similar cell/cluster types that cannot be easily resolved.

## ACKNOWLEDGMENTS

We thank the GenomEast platform in Strasbourg, France. We are grateful to Samah Rekima for technical help on the iBV histology facility and we also thank the iBV PRISM Microscopy facility for their regular support. This work was funded by the French Government (National Research Agency, ANR) through the ‘Investments for the Future’ programs IDEX UCAJedi ANR-15-IDEX-01, by the “Fondation pour la Recherche Médicale (Equipe FRM2020)” (#EQU202003010222), “Fondation de France” (#00123416), ERA-NET Neuron grant (Brain4Sight) (ANR-21-NEU2-0003-03) grants to M.S., by the ‘France Génomique’ consortium (#ANR-10-INBS-0009) to M.J and by INSERM funding (“Dotation exceptionnelle prise de fonctions”) to M.B.

## AUTHOR CONTRIBUTIONS

Conceptualization, M.B. and M.S.; Methodology, M.B., M.S. and M.J.; Investigation, M.B., G.M., A.L. and M.J.; Writing – Original Draft, M.B., M.J. and M.S.; Writing – Review & Editing, M.B. and M.S.; Visualization, M.B.; Supervision, M.S.; Funding Acquisition, M.B. and M.S.

## CONFLICT OF INTEREST STATEMENT

The authors declare that they have no conflict of interest.

## Notes

### Competing Interest Statement

The authors have declared no competing interest.

