## Supplementary Information for "FGF8-mediated gene regulation affects regional identity in human cerebral organoids"

### *SUPPLEMENTAL FIGURES AND LEGENDS*

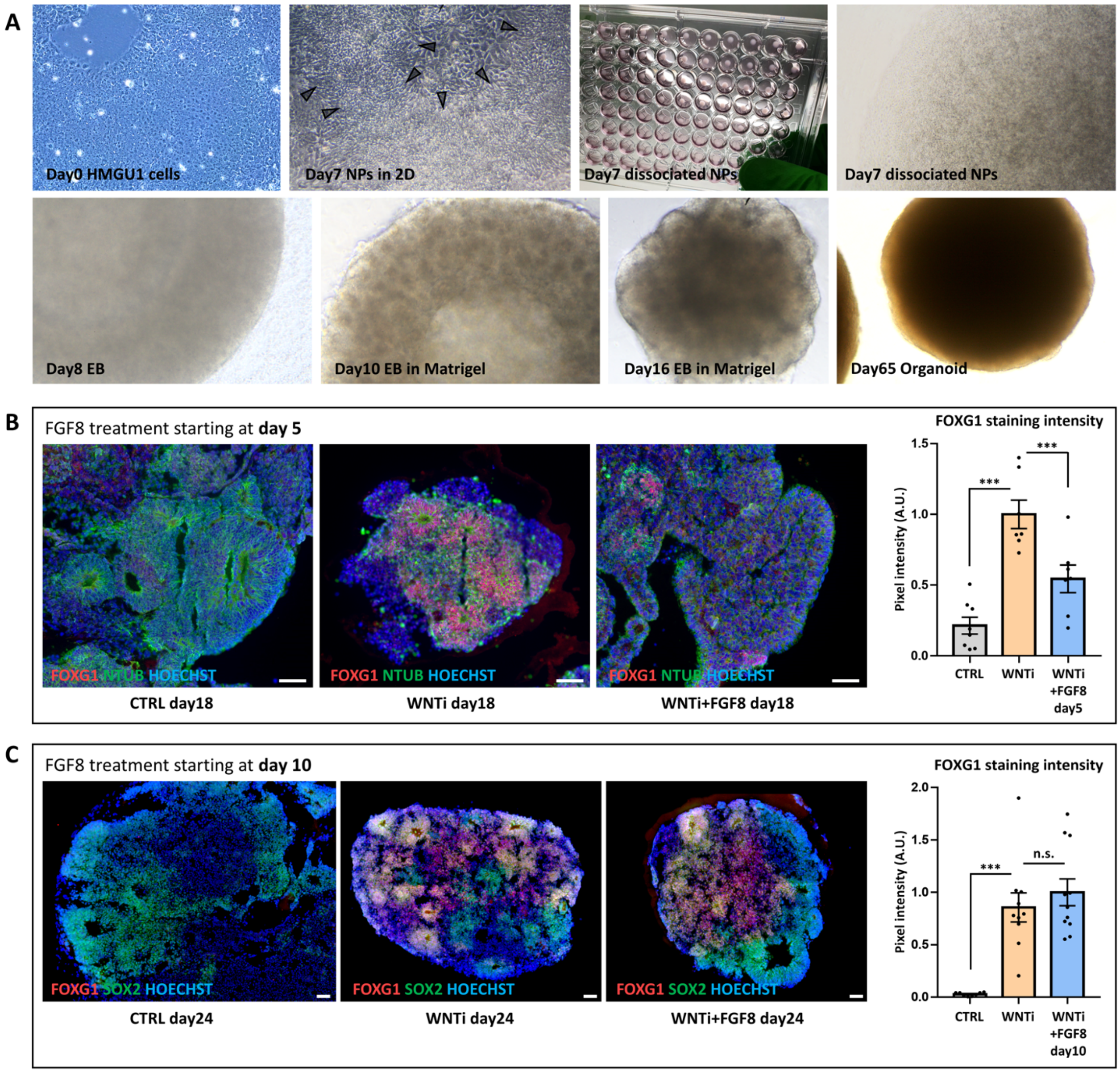


***Figure S1. Hybrid 2D/3D protocol and effect of early FGF8 treatment on FOXG1 expression in human organoids. A)*** *Brightfield microscope images showing, from top left to bottom right: undifferentiated and confluent HMGU1 cells at the starting day of the neural differentiation protocol; day7 neural progenitors organized in groups of radially oriented cells (arrowheads) and surrounded by non-neural cells; dissociated NPs at day7, spinned at the bottom of 96-well plates; day7 dissociated NPs at the bottom of a single well; embryoid body (EB) at day8, 24 hours after dissociation; day10 EB included in Matrigel; EB at day16, with some rosettes and neural epithelia becoming visible on the borders; day65 organoid.* ***B,C)*** *Panels showing FOXG1 (red) and NTUB (green) immunostaining in control (CTRL), WNT inhibited (WNTi) and FGF8 treated (WNTi+FGF8) organoids, as indicated. When added at day5 (Panel A), FGF8 partially inhibited FOXG1 expression, suggesting interference with induction of telencephalic identity (pixel intensity in graph). FGF8 treatment starting at a later time point (day8; Panel B) allowed for efficient FOXG1 induction (pixel intensity in graph). Scale bars: 50 µm.*

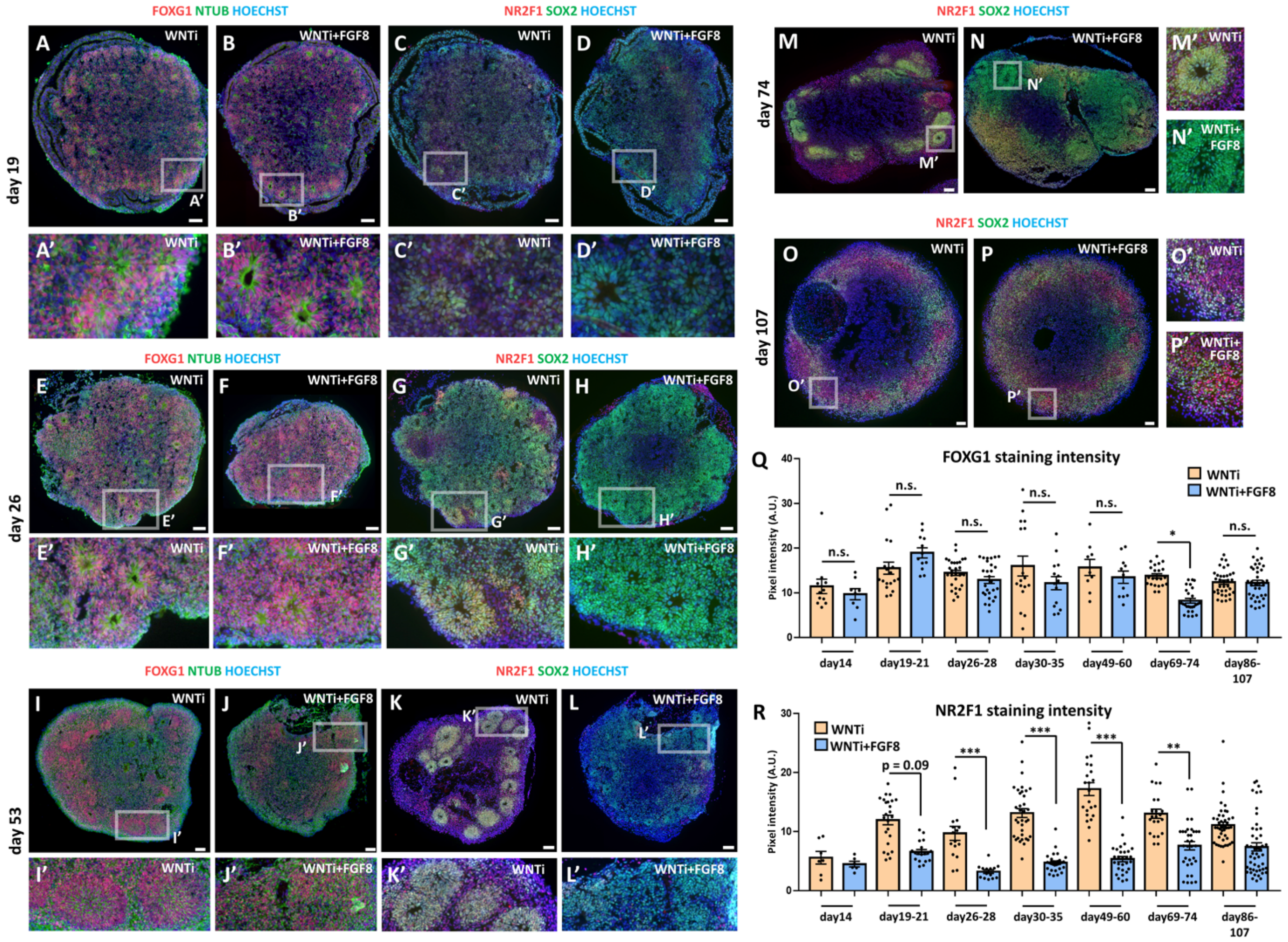


***Figure S2. FGF8-mediated control of NR2F1 level in FOXG1+ telencephalic organoids. A-H')*** *FOXG1 and NTUB (red and green, respectively, in A-B' and in E-F') and NR2F1 and SOX2 (red and green, respectively, in C-D' and in G-H') immunostainings on day19 or on day26 WNTi and WNTi+FGF8 organoids, as indicated. FGF8 treatment did not significantly interfere with FOXG1 expression (A-B',E-F'), while NR2F1 (which started to be detectable at low levels by immunostaining at day19 in WNTi organoids) was down-regulated by FGF8 (compare C,C' with D,D' or G,G' with H,H').* ***I-L')*** *FOXG1 and NTUB (red and green, respectively, in I-J') and NR2F1 and SOX2 (red and green, respectively, in K-L') immunostainings on day53 WNTi and WNTi+FGF8 organoids, as indicated.* ***M-P')*** *NR2F1 (red) and SOX2 (green) immunostaining in day74 (M-N') and in day107 (O-P') WNTi and WNTi+FGF8 organoids, showing that NR2F1 is still efficiently modulated by FGF8 20-25 days after end of the treatment (N,N') but is gradually upregulated back to control levels in long term cultured organoids (day107; P-P').* ***Q)*** *Graph shows pixel intensity quantification of FOXG1 levels after immunostaining of WNTi and WNTi+FGF8 organoids at different time points, as indicated.* ***R)*** *Graph shows pixel intensity quantification of NR2F1 immunostaining in WNTi and WNTi+FGF8 organoids at different time points, as indicated. NR2F1 level was efficiently downregulated by FGF8 treatment from day26 to day74, while it raised back at later time points. Scale bars: 100 µm.*

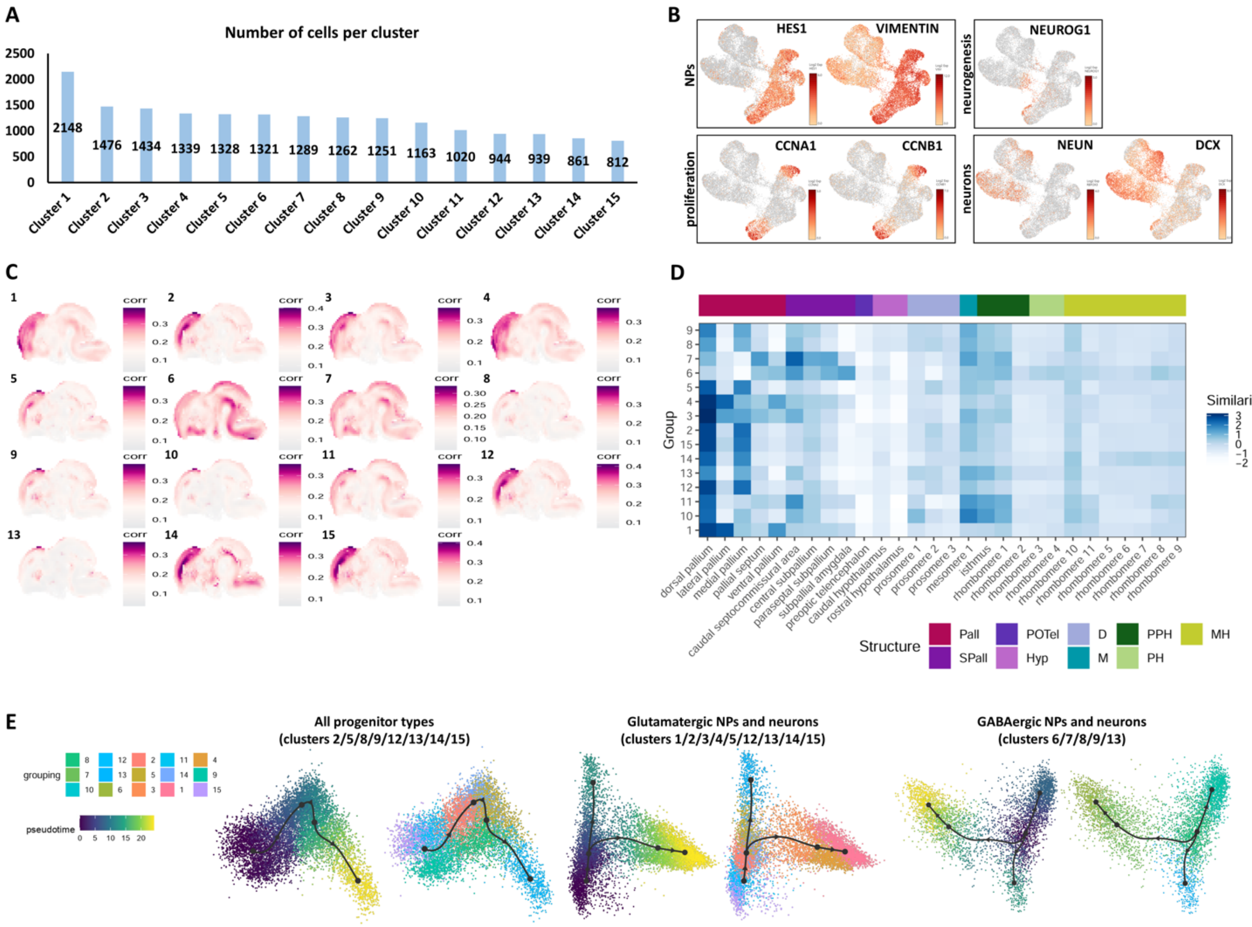

***Figure S3. Single-cell RNA sequencing (scRNAseq) and VoxHunt similarity map analysis of two-months old telencephalic organoids. A)*** *Number of cells per cluster after scRNAseq of day69 organoids.* ***B)*** *Expression level of known markers identifying neural progenitor cells (NPs; VIMENTIN and HES1), proliferating progenitors (CCNA1 and CCNB1), differentiating neurons (NEUROG1) and post-mitotic neurons (DCX and NEUN). Additional markers are shown in Figure 3C.* ***C)*** *VoxHunt similarity map showing similarity correlation index (white to violet) of each cluster to reference databases of mouse regional brain atlases.* ***D)*** *VoxHunt heatmap of similarity score showing the similarity degree (blue color code) between the expression profile of clusters (lines) and distinct regions of the mouse embryonic brain (columns). Note the high similarity between clusters 1/2/3/4/5/12/14/15 and dorsal-anterior pallium (neocortex). Clusters 6 and 7, on the contrary, show high similarity to ventral subpallial regions (ganglionic eminences). Legend on the right shows the brain area associated to each color. Pall: pallium; Spall: Subpallium; POTel: Preoptic telencephalon; Hyp: hypothalamus; D: diencephalon; M: mesencephalon; PPH: Prepontine hindbrain; PH: Pontine hindbrain; MH: Medullary hindbrain.* ***E)*** *Trajectory analysis evaluating the most probable developmental trajectory linking all cell progenitor clusters (left), glutamatergic NPs and neurons (center) or GABAergic NPs and neurons (right). NP clusters can either convert into one another (suggesting plasticity to switch between dorsal and ventral telencephalic identity), either differentiate into post-mitotic neurons. In all cases, progenitors can also enter in a developmental “bottleneck” represented by cluster 13 (CRYAB+ truncated aRGs).*

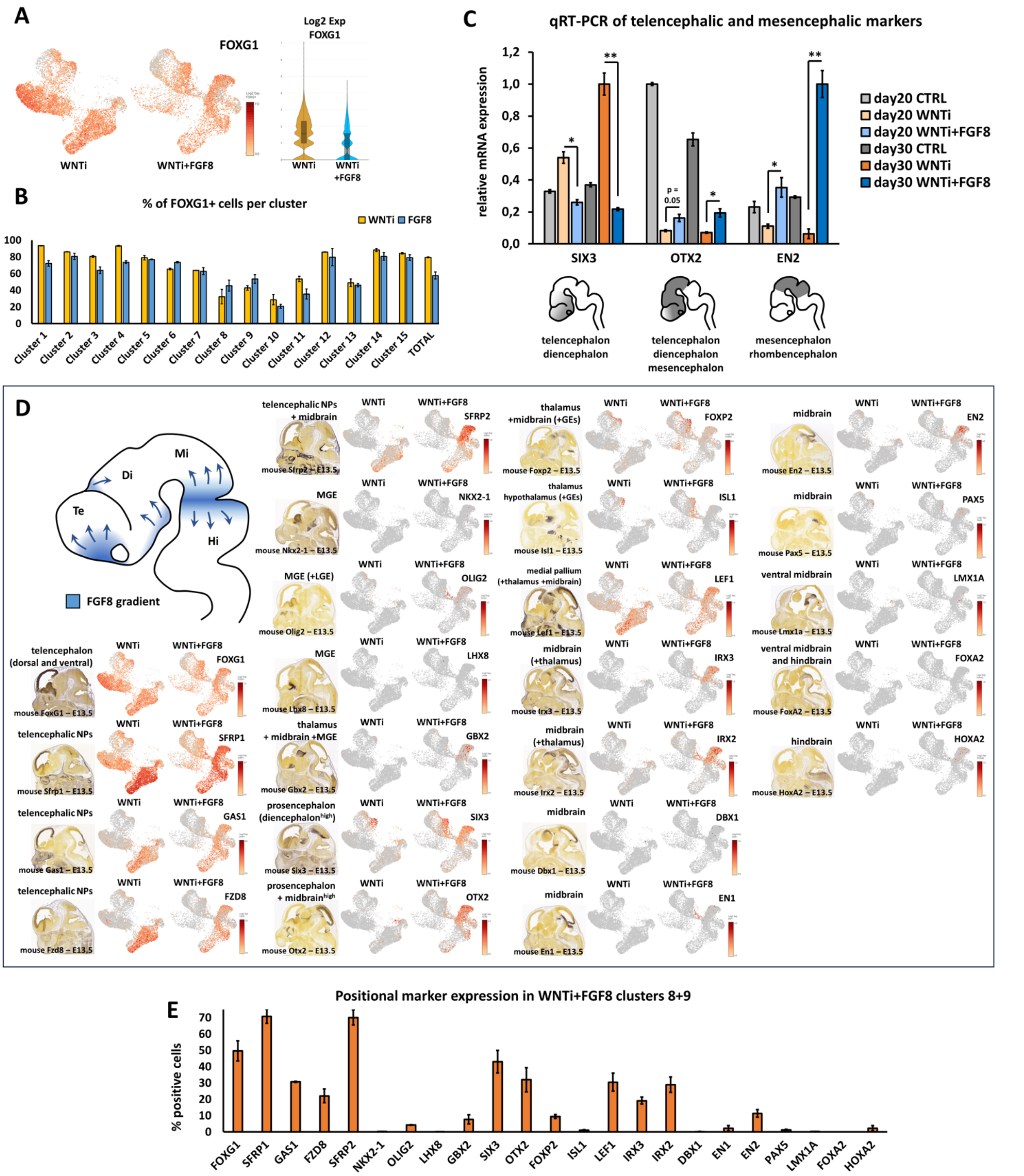


***Figure S4. FGF8-dependent induction of diencephalic and mesencephalic markers in telencephalic organoids. A,B)*** *Expression level and percentages of positive cells per cluster for FOXG1 in control (WNTi) and treated (WNTi+FGF8) organoids, as indicated. Despite a partial reduction of FOXG1 average expression (A), most of the cells in different clusters still expressed FOXG1 after FGF8 treatment (B).* ***C)*** *Real time RT-PCR quantification of telencephalic/diencephalic (SIX3), prosencephalic/midbrain (OTX2) and midbrain/hindbrain (EN2) markers in day20 or day30 control (CTRL), WNT inhibited (WNTi) or FGF8-treated (WNTi+FGF8) organoids, as indicated. FGF8 induced loss of SIX3 expression and increase of diencephalon/midbrain markers (OTX2, EN2), suggesting that long term FGF8 treatment can induce additional -more posterior- regional identities together with FOXG1+ cells.* ***D)*** *At the top left of the panel, an early fetal brain scheme shows FGF8 sources (blue) in the anterior telencephalon, in the diencephalon and at the midbrain/hindbrain border, and their presumable diffusing gradients (arrows). The panel shows the expression levels of distinct brain regional markers in UMAP projections of WNTi or WNTi+FGF8 single cell RNA sequencing samples, as indicated. Mouse brain images at*

*embryonic day (E) 13.5 obtained from the Allen Brain Atlas show the main domains where these markers are found.* ***E)*** *Graph showing the percentage of cells expressing different markers (indicated on x-axis) in clusters 8 and 9 of WNTi+FGF8 organoids. Clusters 8/9 showed high percentage of cells expressing telencephalic (FOXG1, SFRP1, GAS1, SFRP2) and diencephalic/midbrain markers (SIX3, OTX2, LEF1, IRX3, IRX2, EN2). Very few or no cells expressed markers for medial ganglionic eminence (GBX2, LHX8, NKX2-1, OLIG2), for ventral midbrain (FOXP2, ISL1, LMX1A, FOXA2) or for more posterior regions (HOXA2).*

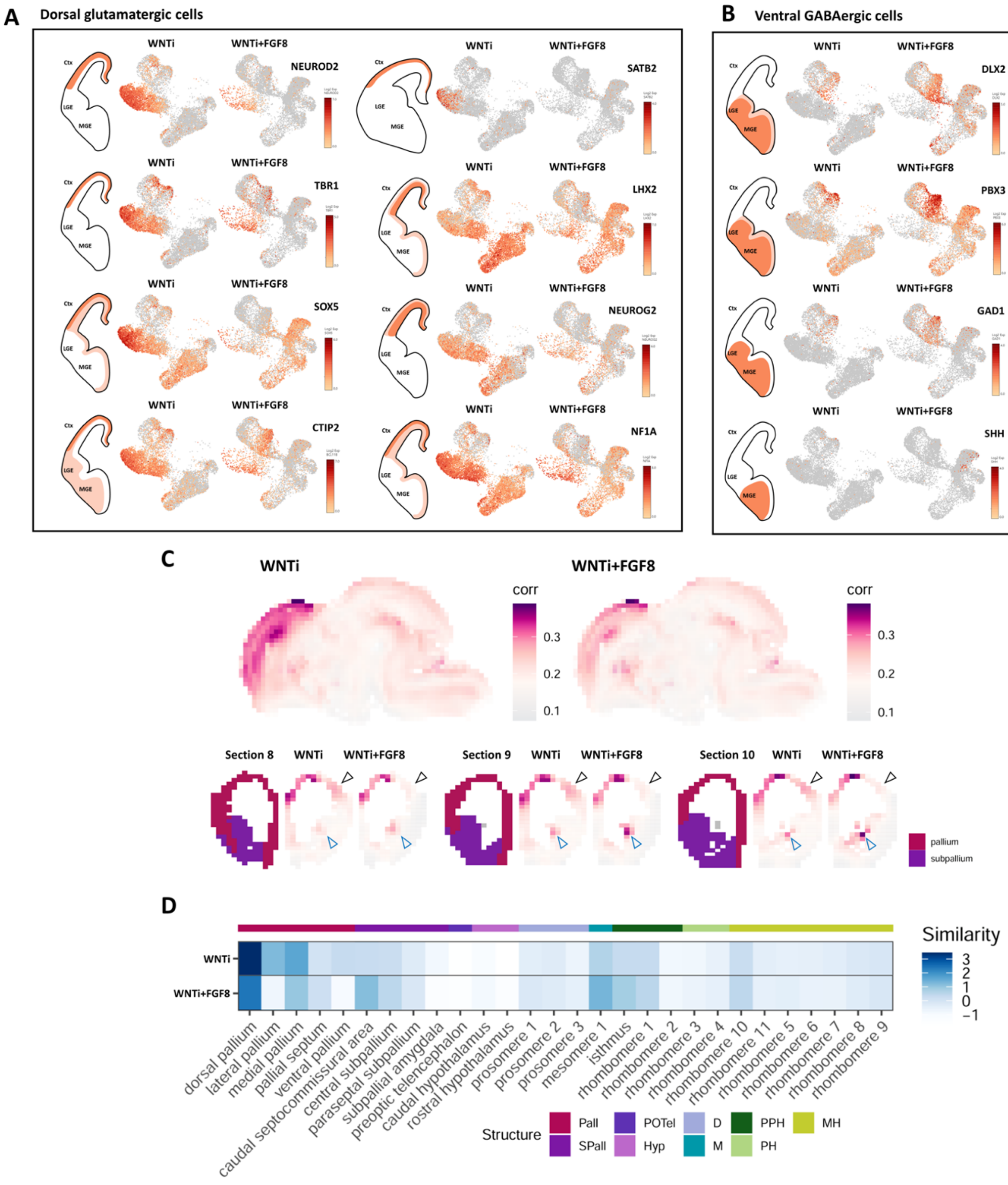


***Figure S5. Changes in cellular composition and glutamatergic/GABAergic identity upon FGF8 treatment. A,B)*** *Expression level of known markers identifying dorsal glutamatergic NPs and neurons (panel A; NEUROD2, TBR1, SOX5, CTIP2, SATB2, LHX2, NEUROG2 and NF1A) or ventral GABAergic NPs and neurons (panel B; DLX2, PBX3, SHH and GAD1). FGF8 treatment caused an increase of GABAergic cells expressing markers of the LGE, at the expense of cells expressing glutamatergic genes. Notably, MGE markers such as NKX2-1, SHH and LHX8 were not significantly expressed in FGF8-treated samples.* ***C)*** *VoxHunt similarity map showing similarity correlation index (white to violet color code) of WNTi (left) or WNTi+FGF8 (right) organoids to reference databases of mouse regional brain atlases. For this analysis, all WNTi cells and all WNTi+FGF8 cells are evaluated. FGF8-treated organoids showed lower similarity to dorso-lateral pallium (black arrowheads) and increased similarity to subpallial areas (blue arrowheads), compared to control samples.* ***D)*** *VoxHunt heatmap of similarity score showing the similarity degree (blue color code) between the expression profile of organoid samples (WNTi -upper line- and WNTi+FGF8 -lower line-) and distinct regions of the human fetal brain (columns). FGF8-treated organoids showed lower similarity to dorsal pallium and increased similarity to subpallial or to more posterior (midbrain) regions, compared to control samples. Pall: pallium; Spall: Subpallium; POTel: Preoptic telencephalon; Hyp: hypothalamus; D: diencephalon; M: mesencephalon; PPH: Prepontine hindbrain; PH: Pontine hindbrain; MH: Medullary hindbrain.*

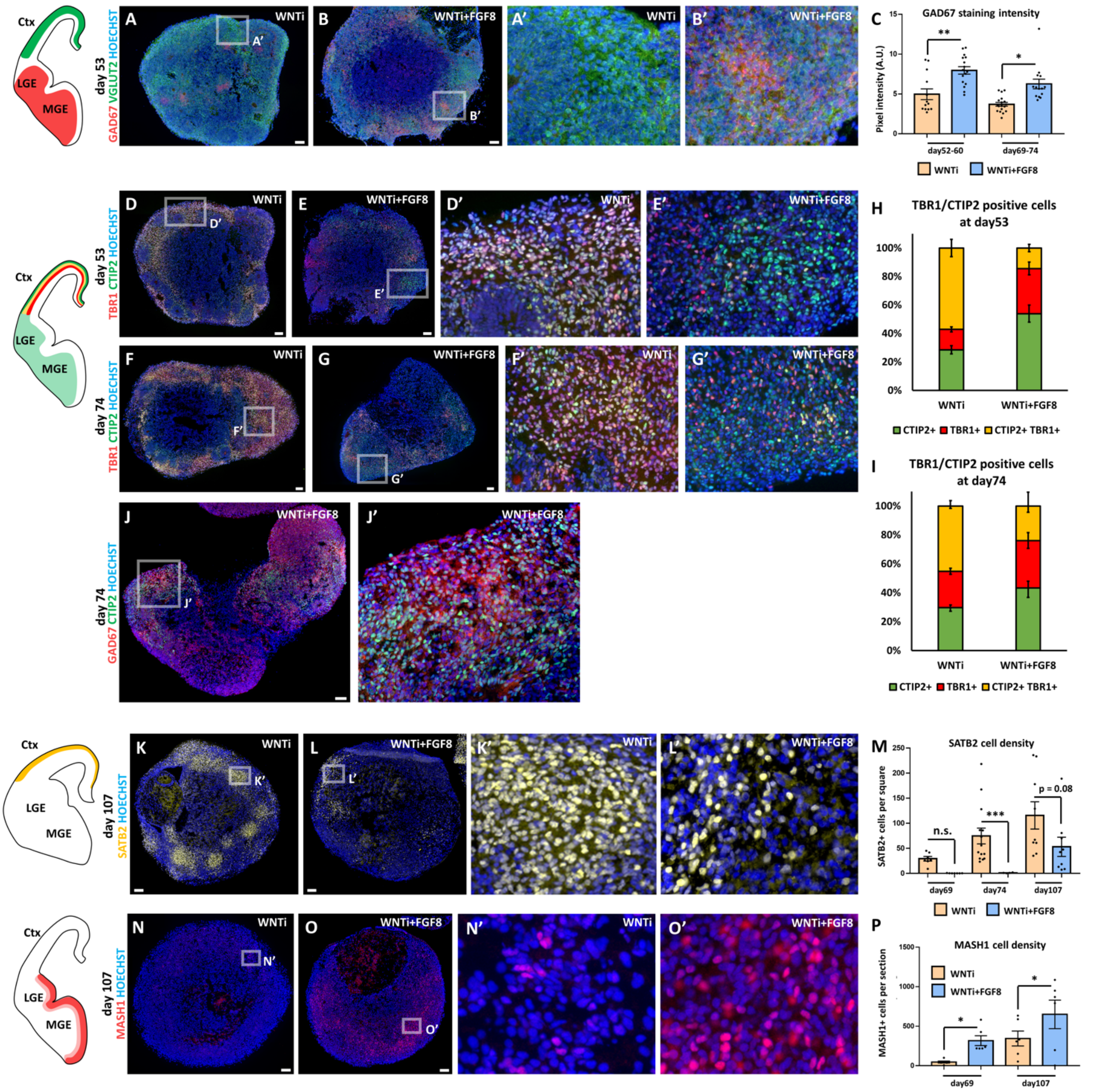

***Figure S6. Immunostaining analysis of glutamatergic and GABAergic cellular composition of control and FGF8-treated telencephalic organoids. A-C)*** *GAD67 (red) and VGLUT2 (green) immunostaining in day53 control (WNTi) and treated (WNTi+FGF8) organoids, as indicated. The distribution of these markers in vivo is shown in the brain scheme on the left, while graph in C displays GAD67 pixel intensity quantification at day52-60 and at day69-74.* ***D-I)*** *TBR1 (red) and CTIP2 (green) immunostaining in day53 (D-E') or day74 (F-G') control (WNTi) and treated (WNTi+FGF8) organoids, as indicated. The distribution of these markers in vivo is shown in the brain scheme on the left. Note that TBR1 and CTIP2 can be co-expressed dorsally (in cortical layers) during development; however, while TBR1 is exclusively dorsal, CTIP2 is also expressed in the ventral telencephalon. Graphs show the percentage of TBR1+ (red), CTIP2+ (green) or double TBR1+/CTIP2+ (yellow) cells in WNTi or WNTi+FGF8 organoids at different steps (day53 in H and day74 in I), as indicated.* ***J,J')*** *GAD67 (red) and CTIP2 (green) immunostaining in day74 WNTi+FGF8 organoids, showing double CTIP2+ GAD67+ cells.* ***K-M)*** *SATB2 (yellow) immunostaining in day107 control (WNTi) and treated (WNTi+FGF8) organoids, as indicated. The distribution of SATB2+ neurons in vivo is shown in the brain scheme on the left, while graph in M displays cell density at day69, day74 or day107. SATB2 cell density was measured as n° of SATB2+ neurons per 150X150µm square.* ***N-P)*** *MASH1 (red) immunostaining in day107 control (WNTi) and treated (WNTi+FGF8) organoids, as indicated. Quantification of the cell density (n° of cells per organoid section) of MASH1+ progenitors at day69 or day107 is shown in the graph in P. Scale bars: 100 µm.*

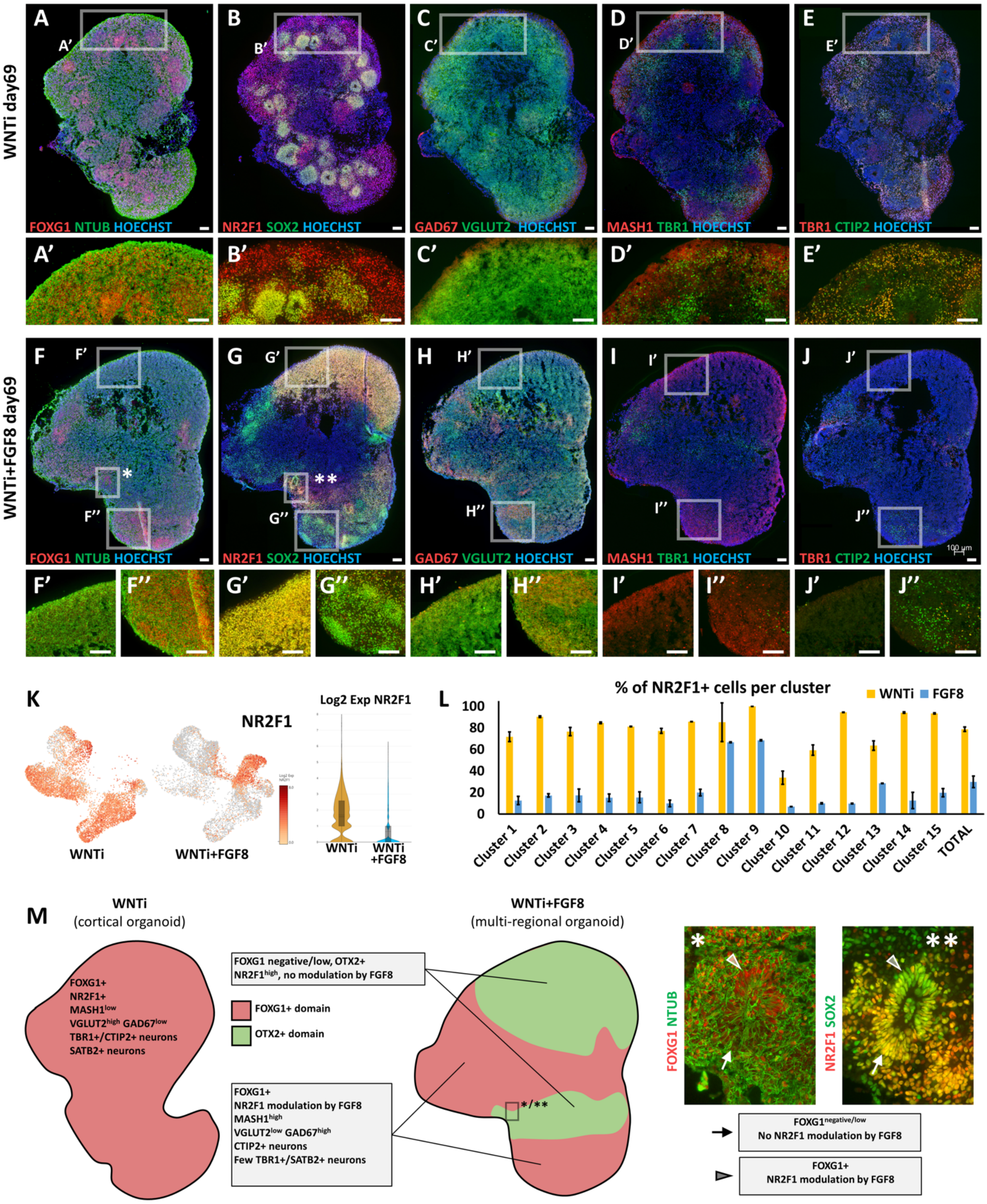


***Figure S7. Regional identity-dependent effect of FGF8 treatment on D/V identity and target gene modulation. A-E')*** *Immunostaining of key neural and regional markers on day69 WNTi control organoids; FOXG1 (red in A,A'), NR2F1 (red in B,B') and SOX2+ NPs (green in B,B'). Glutamatergic/cortical markers VGLUT2 (green in C,C'), TBR1 (green in D,D' or red in E,E') and CTIP2 (green in E,E') are abundant, indicating neocortical identity.* ***F-J'')*** *Immunostaining of the same key neural and regional markers showed in A-E', but on sections of day69 WNTi+FGF8 treated organoids. Insets show magnification from a FOXG1-negative non-telencephalic region (F',G',H',I',J') or from a FOXG1+ telencephalic region (F'',G'',H'',I'',J''). Notably, NR2F1 (red in G-G'') showed high intensity, i.e. no response to FGF8 treatment, in FOXG1-negative non-telencephalic areas. Only FOXG1+ telencephalic domains showed FGF8-mediated reduction of NR2F1 levels (G,G''), associated with induction of ventral markers GAD67 (red in H,H'') and MASH1 (red in I,I'') at the expense of dorsal*

*cortical markers TBR1 (green in I-I''). No dorsal cortical markers such as TBR1 and CTIP2 were detectable in FOXG1-negative regions (I',J'), showing that efficient acquisition of anterior identity is necessary for production of cortical neurons in organoids. Insets indicated by asterisks in F (*) and G (**) are showed at high magnification in M. Scale bars: 100 µm.* ***K,L)*** *Expression level and percentages of positive cells per cluster for NR2F1 in control (WNTi) and treated (WNTi+FGF8) organoids, as indicated. NR2F1 average level was greatly reduced by FGF8 (K); however, clusters 8 and 9 did not respond to FGF8 treatment by downregulating NR2F1 (L) and still contained >60% NR2F1+ cells.* ***M)*** *Schematic representation of distinct domains in control (WNTi; left) and FGF8-treated (WNTi+FGF8; right) organoids. FOXG1+ telencephalic WNTi organoids (left) show abundant levels of glutamatergic marker VGLUT2 and abundant SATB2+ and TBR1/CTIP2 double positive cortical neurons. Upon FGF8 treatment, WNTi+FGF8 organoids (right) develop co-existing FOXG1+ telencephalic domains (light red areas), in which ventral GABAergic markers GAD67 and MASH1 are upregulated at the expense of dorsal genes, and FOXG1-negative/OTX2+ diencephalic/midbrain domains (light green areas). Immunostaining magnifications on the right (magnification * and ** from images in Suppl. Figure 7) show a neural rosette located at the border between a FOXG1+ domain (upper half of the rosette; arrowheads) and an adjacent FOXG1-low domain (lower half; arrows), to highlight that NR2F1 was down-regulated by FGF8 only in the telencephalic domain (upper half of the rosette), indicating that FGF8 target genes can respond in a different way depending on the regional identity of organoid domains.. Scale bars: 100 µm.*

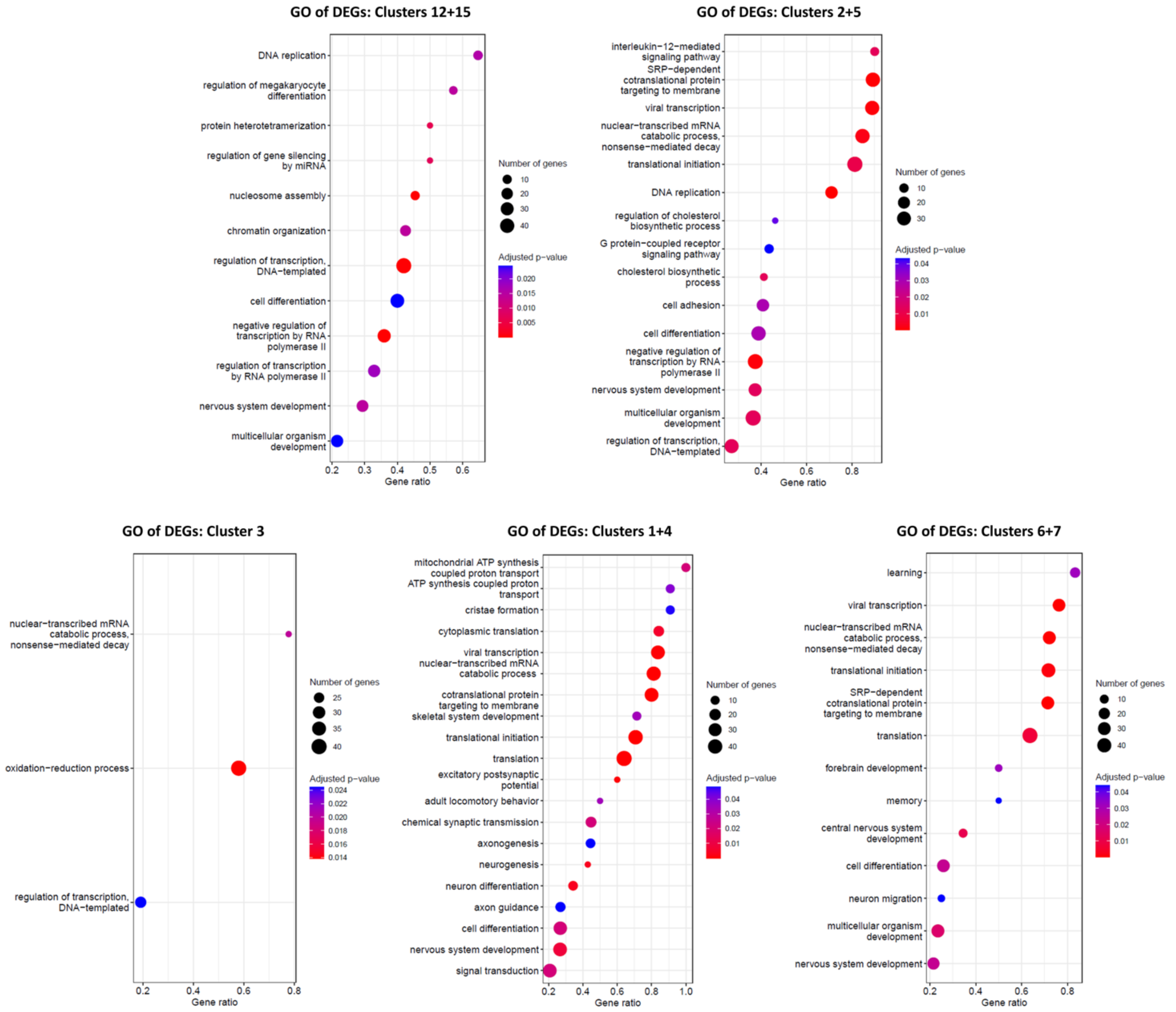


***Figure S8. Enrichment analysis of Gene Ontology (GO) terms in distinct NP or neuronal clusters.*** *Results of enrichment analysis of the biological process GO terms performed with the gene set enrichment analysis (GSEA) method. Each graph shows the results relative to a specific cluster or subset of clusters, as indicated. Red to blue color code corresponds to adjusted p-value, while the circle size indicates the number of genes that belong to a specific GO term listed on the left of each graph.*
